# Comprehensive Transcriptomic and Epigenomic Insights into Environmental Toxicant Exposures: The TaRGET II Resource

**DOI:** 10.1101/2025.07.28.667191

**Authors:** Bo A. Zhang, Benpeng Miao, Shuhua Fu, Cristian Coarfa, Ravindra Kumar, Prashant Kumar Kuntala, Bongsoo Park, Sandra l. Grimm, Rahul Jangid, Justin A. Colacino, Laurie K. Svoboda, Wanqing Shao, Xiaoyun Xing, Daofeng Li, Shaopeng Liu, Robert B. Hamanaka, Claudia Lalancette, Maureen A. Sartor, Christopher Krapp, Gregory E Crawford, Heather B. patisaul, Tim Wiltshire, TaRGET II Consortium, David Aylor, Shyam Biswal, Gökhan M. Mutlu, Sanjay Rajagopalan, Wan-Yee Tang, Ting Wang, Dana C. Dolinoy, Marisa S. Bartolomei, Cheryl Walker

## Abstract

Environmental exposures to toxic chemicals can profoundly alter the transcriptome and epigenome in both humans and animals, contributing to disease development across the lifespan. To elucidate how early-life exposure to toxicants exerts such persistent effects, the *Toxicant Exposures and Responses by Genomic and Epigenomic Regulators of Transcription II (TaRGET II)* Consortium generated a landmark resource comprising 2,564 epigenomes and 1,043 transcriptomes from longitudinal studies in mice. All data are publicly available through the <u>TaRGET II data portal</u> and the WashU Epigenome Browser. This resource from target (liver, brain, lung, heart) and surrogate (blood) tissues at weaning (3 weeks) and two adult time-points (5 and 10 months) characterized the molecular response to arsenic (As), lead (Pb), bisphenol-A (BPA), di-2-ethylhexyl phthalate(DEHP), tributyltin (TBT), tetrachlorodibenzo-p-dioxin (TCDD), and particulate matter with a diameter of <2.5μm (PM_2.5_). The findings revealed persistent, toxicant-specific, sex-dependent epigenomic and transcriptomic perturbations, resulting in disrupted expression of 14,908 genes, altered chromatin accessibility at 87,409 regulatory elements, DNA methylation changes at 113,186 genomic regions, and chromatin state switching of histone modifications. The resulting high-resolution map of how environmental exposures reprogram the epigenome and transcriptome is broadly accessible via *ToxiTaRGET* database, offering unparalleled opportunities for the scientific community to investigate the molecular underpinnings of environmental toxicant exposures and their contributions to disease pathogenesis.

## Main

Environmental exposures, especially to toxic chemicals, can substantially shape the epigenome in both humans and animals, and significantly impact the processes of development, aging, and disease pathogenesis^1–3^. As estimated in 2022, environmental factors are responsible for about 9 million deaths per year, corresponding to one in six deaths worldwide^4^. Exposure to metals and metalloids, such as arsenic or lead, is associated with many diseases, such as cancer, cardiovascular disease, neurological disorders, and autoimmune diseases^5–7^. Endocrine-disrupting chemicals (EDCs) are associated with human reproductive and metabolic disorders, diabetes, and increased risk of cancer^8–11^, while air pollution, especially in the form of particulate matter with a diameter of <2.5μm, is tightly associated with heart disease, respiratory infections, chronic lung disease, insulin resistance^12^, and other illnesses^13–16^. Thus, it is critical to understand at the molecular level the response to toxic environmental exposures, which eventually facilitates precision environmental health solutions to improve health and lessen the burden of environmental disease^17–20^.

The epigenome is both mitotically heritable and dynamic, exhibiting context-specific changes all across the life course^21–25^. These changes contribute to the numerous distinct gene expression programs that mediate biological functions in response to complex developmental, as well as environmental cues^26–28^. To systematically elucidate the epigenomic and transcriptomic responses to toxicant exposures and underlying molecular mechanisms, the *Toxicant Exposures and Responses by Genomic and Epigenomic Regulators of Transcription II (TaRGET II)* Consortium^29^ was established. The TaRGET II project systematically investigated transcriptomic and epigenomic responses in multiple tissues following early-life exposure to a diverse array of environmental toxicants. Exposures were administered from preconception through lactation via maternal oral and inhalation routes (see Methods). Subsequent profiling was conducted at three stages: weaning (∼3 weeks), young (5 months), and later (10 months) adulthood [**Figure 1**]. *TaRGET II* has now constructed the most comprehensive catalog to date of molecular perturbations induced by environmentally relevant exposures. These include arsenic (As), lead (Pb), EDCs bisphenol A (BPA), tributyltin (TBT), di-2-ethylhexyl phthalate (DEHP), tetrachlorodibenzo-p-dioxin (TCDD), and particulate matter <2.5 μm (PM_2.5_) with distinct components [**Figure 1**, **Table 1**]. This catalog provides a robust foundation for understanding epigenome-environment interactions that shape health and disease trajectories. TaRGET II data have been assembled into a publicly accessible repository at the TaRGET II data portal (https://dcc.targetepigenomics.org/) and an accompanying database (https://toxitarget.com/ ), creating a valuable scientific and biomedical research resource.

**Figure 1.**
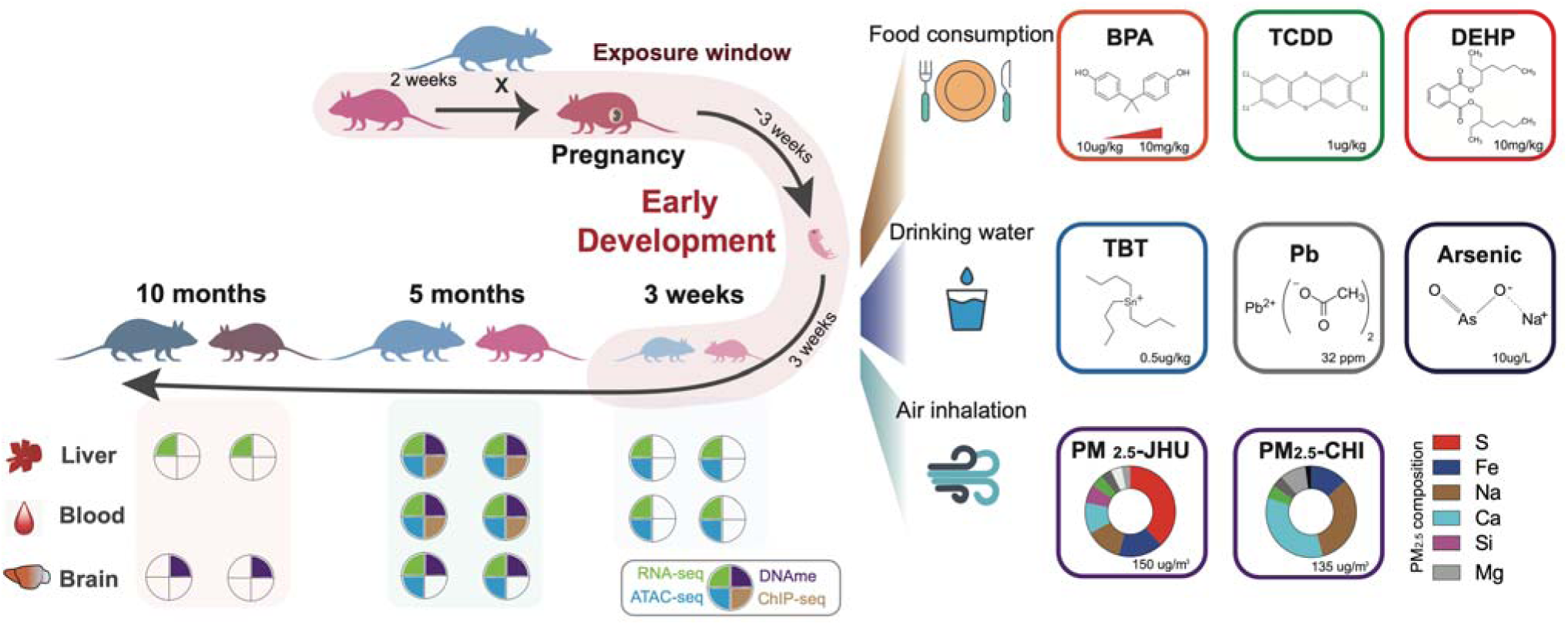
Female mice were exposed to the environmental toxicants shown (Methods) through food, water, and air from 2 weeks pre-conception through pregnancy until weaning at 3 weeks of age. Exposed and matched vehicle-control offspring were raised to 10 months of age without exposure. Tissues were collected and assayed at 3 weeks, 5 months, and 10 months, with the litter used for determining the numbers in subsequent analysis.

**Table 1.** Environmental toxicants included in TaRGET II consortium.

| Toxicant | Exposure type | Dose | Health threats | Impacted population |
| --- | --- | --- | --- | --- |
| **Arsenic** | Drinking water | 10 µg/L/q.o.d | neurodevelopmental toxicity, metabolic effects, and cancer<sup>30-32</sup> | >226 million with unsafe concentrations exposure<sup>33</sup> |
| **BPA** | Chow food | 50 µg/kg, 50 mg/kg | hormonal disruptions, metabolic and reproductive effects, cardiovascular disease, obesity<sup>34-36</sup> | >1µg/L in the urine of > 90% of population<sup>37</sup> |
| **DEHP** | Chow food | 25 mg/kg | metabolic and reproductive effects<sup>38,39</sup> | detectable levels in the urine of >75% of the population<sup>40</sup> |
| **Pb** | Drinking water | 32 ppm | cognitive deficits, developmental delays, cardiovascular issues, kidney damage, and cancer<sup>41-43</sup> | ~800 million children worldwide exposed to high levels of lead<sup>44</sup> |
| **PM<sub>2.5</sub>** | Air inhalation with concentrated PM<sub>2.5</sub> | 135 µg/m<sup>3</sup>(CHI) , 150 µg/m<sup>3</sup>(JHU), 40 hours/week | respiratory and cardiovascular diseases, strokes, lung cancer, and complications from chronic conditions<sup>45-47</sup> | >50% world population live with PM air pollution (PM<sub>2.5</sub>>35µg/m<sup>3</sup>)<sup>48</sup> |
| **TBT** | Drinking water | 0.5 mg/kg | endocrine disorders, metabolic and reproductive effects, cancer<sup>49-51</sup> | population living with contaminated seawater and seafood<sup>52-54</sup> |
| **TCDD** | Oral gavage | 1 µg/kg | hepatotoxicity, carcinogenesis, reproductive disorders, neurodevelopmental toxicity<sup>55-57</sup> | accumulated with ~46pg daily intake from food<sup>58</sup> |

To characterize the molecular changes caused by each environmental toxicant, we used multiple assays, including Assay for Transposase-Accessible Chromatin sequencing (ATAC-seq), Whole Genome Bisulfite Sequencing (WGBS), and Infinium Mouse Methylation BeadChip, Chromatin Immunoprecipitation followed by sequencing (ChIP-seq), and RNA sequencing (RNA-seq). These profiles were generated from individual mice without pooling. The resulting TaRGET II dataset contains 2,564 epigenomes and 1,043 transcriptomes from both target tissues (liver, brain, lung, heart) and surrogate tissue (blood) for both sexes [**Figure 2a**]. We also integrated these multi-life-stage measurements to understand how these exposures altered the transcriptome and epigenome (chromatin accessibility, DNA methylation, and histone modifications). This first-of-its-kind characterization of the multi-omic molecular dynamics in response to early-life exposures revealed:

- Correlative exposure signatures in target (liver) and surrogate (blood) tissues in response to As, BPA, DEHP, Pb, PM_2.5_, TBT, and TCDD.
- Exposure-specific perturbations in the epigenome and transcriptome that persisted into adulthood.
- Changes in temporal epigenomic and gene expression patterns in adulthood disrupted by early-life exposures.
- Sex-specific responses of the epigenome and transcriptome in response to early-life exposures.

**Figure 2.**
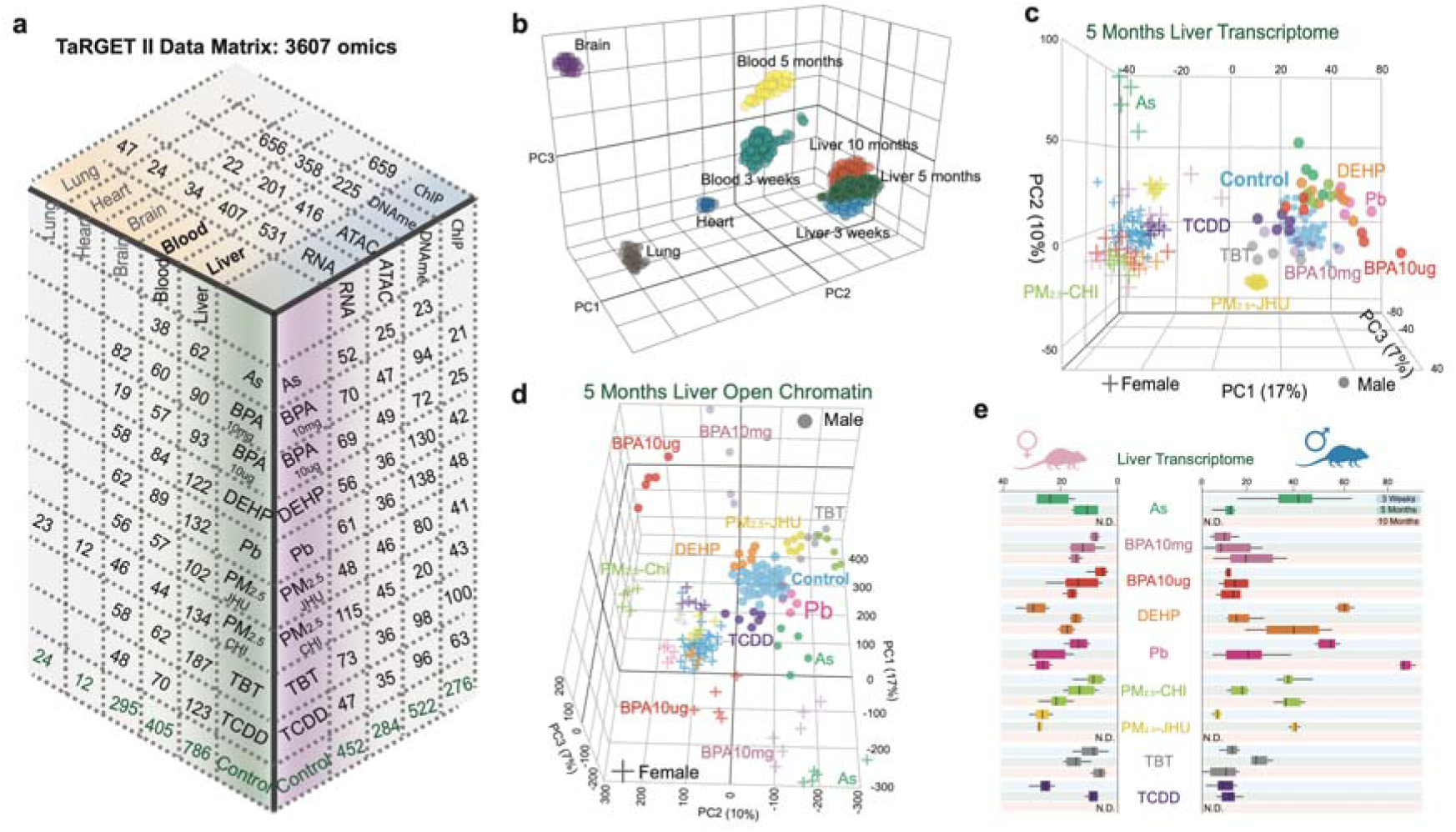
Introduction of TaRGET II dataset. **a**) the number of assayed samples in the data matrix of TaRGET II dataset. 3D PCA illustration of **b)** all tissue transcriptomes from the TaRGET II dataset; **c)** 5-month liver transcriptomes, and **d)** chromatin accessibility, colored by corresponding exposures; **e)** sex-specific Euclidean distance (x-axis) between exposed and age-matched controls (distribution center) in the 3-dimentinal PCA space of the liver transcriptome from all individuals at weaning, young adulthood, and later adulthood.

### Global epigenomic and transcriptomic changes in response to environmental toxicants

TaRGET II generated 3,607 genome-wide omics data sets, including 1,043 RNA-seq data sets, 639 chromatin accessibility data sets, 522 DNA methylation data sets (458 WGBS and 71 RRBS), 744 Infinium Mouse Methylation BeadChip data sets, and 659 profiles for histone modifications on individual mice, providing an unparalleled characterization of the molecular responses to 9 distinct environmental toxicant conditions [**Figure 2a, S-table 1**]. These 2,460 liver and blood datasets were then integrated to systematically explore how early-life exposure interferes with epigenetic programming and gene transcription in mice. We generated normalized gene expression values for RNA-seq^59,60^, genome-wide normalized signal tracks and peaks for ATAC-seq^61–63^, fractional methylation levels for each CpG site^64^, and assessed chromatin states (ChromHMM^65^) using histone modifications. Stringent quality controls were conducted for each data type [**S-table 2**], including calculating inter-replicate correlations and multi-level harmonization of datasets from different production centers [**S-Figure 1,2,3**]. Outlier data sets were flagged and removed from the downstream analysis (see Methods).

Principal Component Analysis (PCA) performed on the integrated datasets revealed that the molecular profiles of target and surrogate tissue samples retained distinct characteristics reflecting their tissue of origin and age [**Figure 2b**]. Additionally, when focusing on a single tissue, such as liver, clear sex-dependent and toxicant-specific responses in the transcriptome [**Figure 2c**] and chromatin accessibility [**Figure 2d**] were evident. For example, chromatin accessibility, measured by differentially accessible regions (DARs), was significantly altered by toxicant exposure at both weaning and young adulthood, depending on sex and exposure [**S-table 3**]. In weanling livers, the greatest increases in accessibility were observed in males exposed to (As), BPA10mg, and TCDD, and in females exposed to TCDD (more than 65% of DARs were more accessible). Reduced accessibility was observed in males responding to PM_2.5_ generated by the University of Chicago consortium (PM_2.5_-CHI), females exposed to BPA10mg and BPA10µg, and TBT (more than 70% of DARs were less accessible). In young adulthood at 5 months, notable increases in chromatin accessibility were only seen in males and females exposed to As. Decreases in accessibility mainly occurred in males and in response to PM_2.5_-CHI, PM_2.5_ from the Johns Hopkins consortium (PM_2.5_- JHU), BPA10mg, BPA10µg, and TBT [**S-table 3**].

Disruption of the transcriptome varied in response to all exposures and was also influenced by both sex and age. For example, in females, the whole transcriptome response to early-life exposure to BPA, Pb, and PM_2.5_-CHI was an increase along with age, compared to age- and sex-matched cont ols [**Figure 2e**]. In contrast, males exposed to BPA10mg did not show an increase in PCA distance as they aged, and Pb-exposed males experienced a decrease in PCA distance between weaning and young adulthood, followed by a much larger increase in later adulthood than females, an intense response not observed with other exposures. Transcriptomic profiling also uncovered notable individual variability in responses to several exposures, most clearly in males exposed to As (3 weeks), Pb (5 months), and DEHP (10 months) [**Figure 2e**]. Interestingly, this variability in transcriptomic disruption was not always reflected in changes in chromatin accessibility, which remained relatively consistent across exposed individuals, even when transcriptomic heterogeneity [**S-Figure 4**] and DNA methylation [**S-Figure 5**] heterogeneity were evident. Therefore, these different methods revealed unique aspects of how the epigenome and transcriptome respond to early-life exposures, highlighting that examining only one “omic” layer might not fully capture the acute and long-term effects of toxicant exposure, and revealed the importance of the Multiomics profiling approach in future biomarker development and understanding molecular mechanisms.

**Figure 3.**
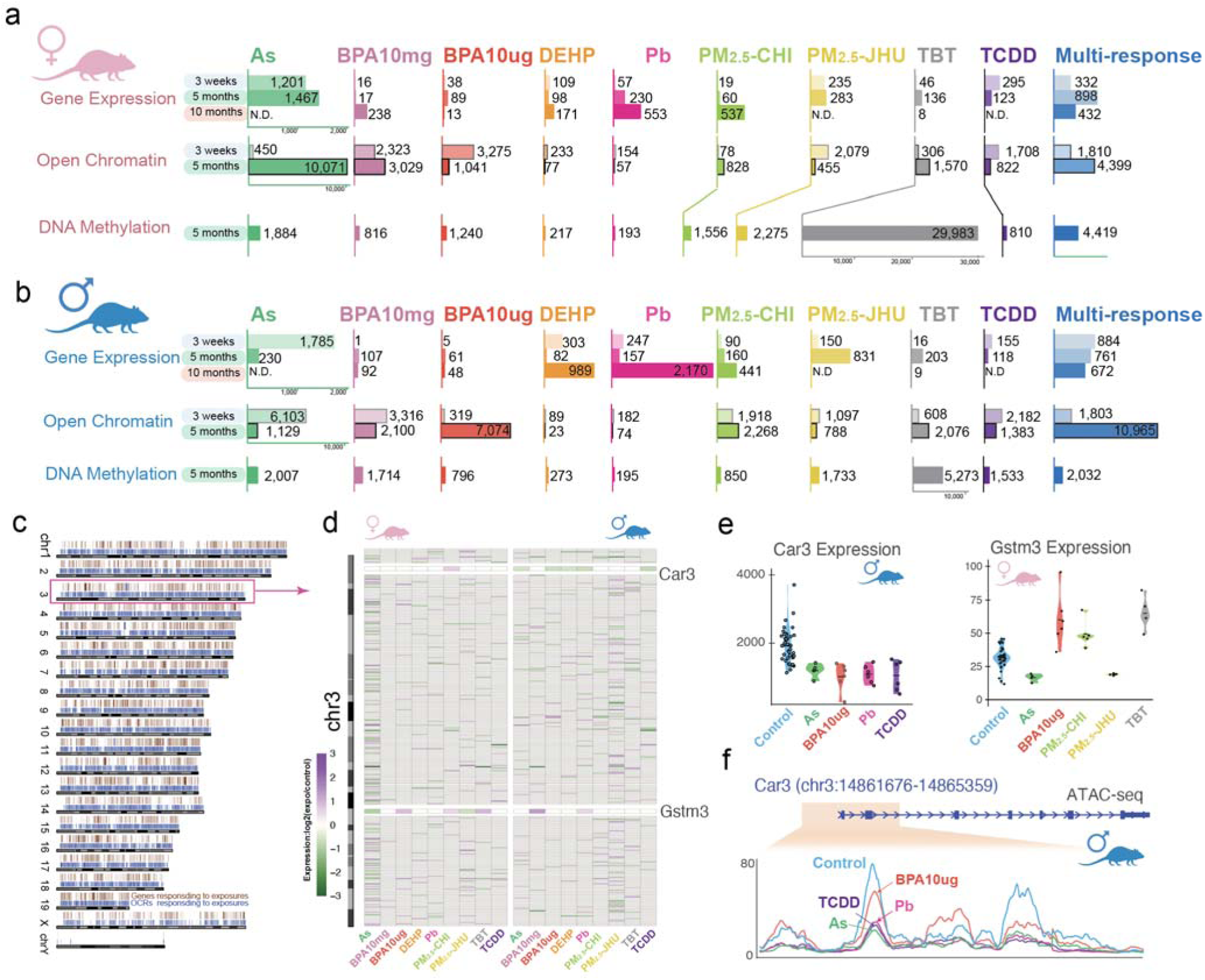
Molecular signatures identified in the TaRGET II dataset as a function of early-life environmental exposures. Numbers of exposure-specific and multi-repsonse differentially expressed genes, accessible regions, and DNA methylated regions for every exposure condition in **a)** female mouse liver and **b)** male mouse liver; **c)** chromosome-scale distribution of all DEGs and DARs; **d)** Differentially expressed chromosome 3 genes (274) at 5 months as a function of toxicant exposures in females and males; **e)** expression levels (RPKM) of *Car3* and *Gstm3*; **f)** averaged open chromatin signals (ATAC-seq) on *Car3* promoter region across exposure types.

**Figure 4.**
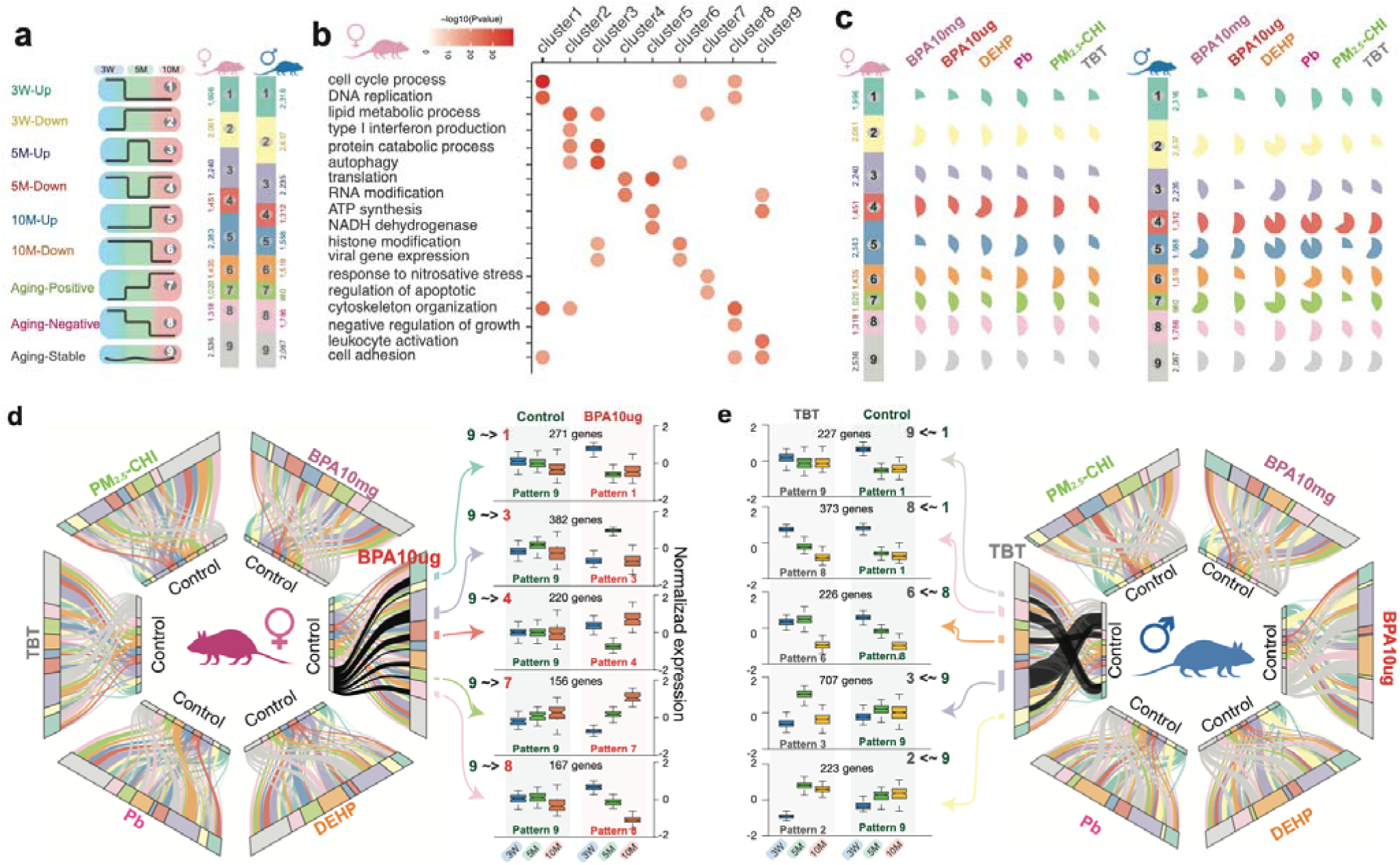
Disrupted temporal expression patterns in mouse liver as a function of early-life environmental exposures. **a)** nine temporal expression patterns across mouse liver development and aging; **b)** enriched biological processes of each temporal expression pattern in female mice; **c)** percentage of genes that changed temporal expression pattern as a function of early-life environmental exposures; **d)** the changes of temporal expression patterns for all genes in females under six environmental exposures (Left). Expression plot of genes that changed temporal patterns from 9 to 1,3,4,7, and 8 under BPA10µg exposure (Right); **e)** expression plot of genes that changed temporal patterns under TBT exposure (Left). The changes of temporal expression patterns for all genes in males as a function of six environmental exposures (Right).

**Figure 5.**
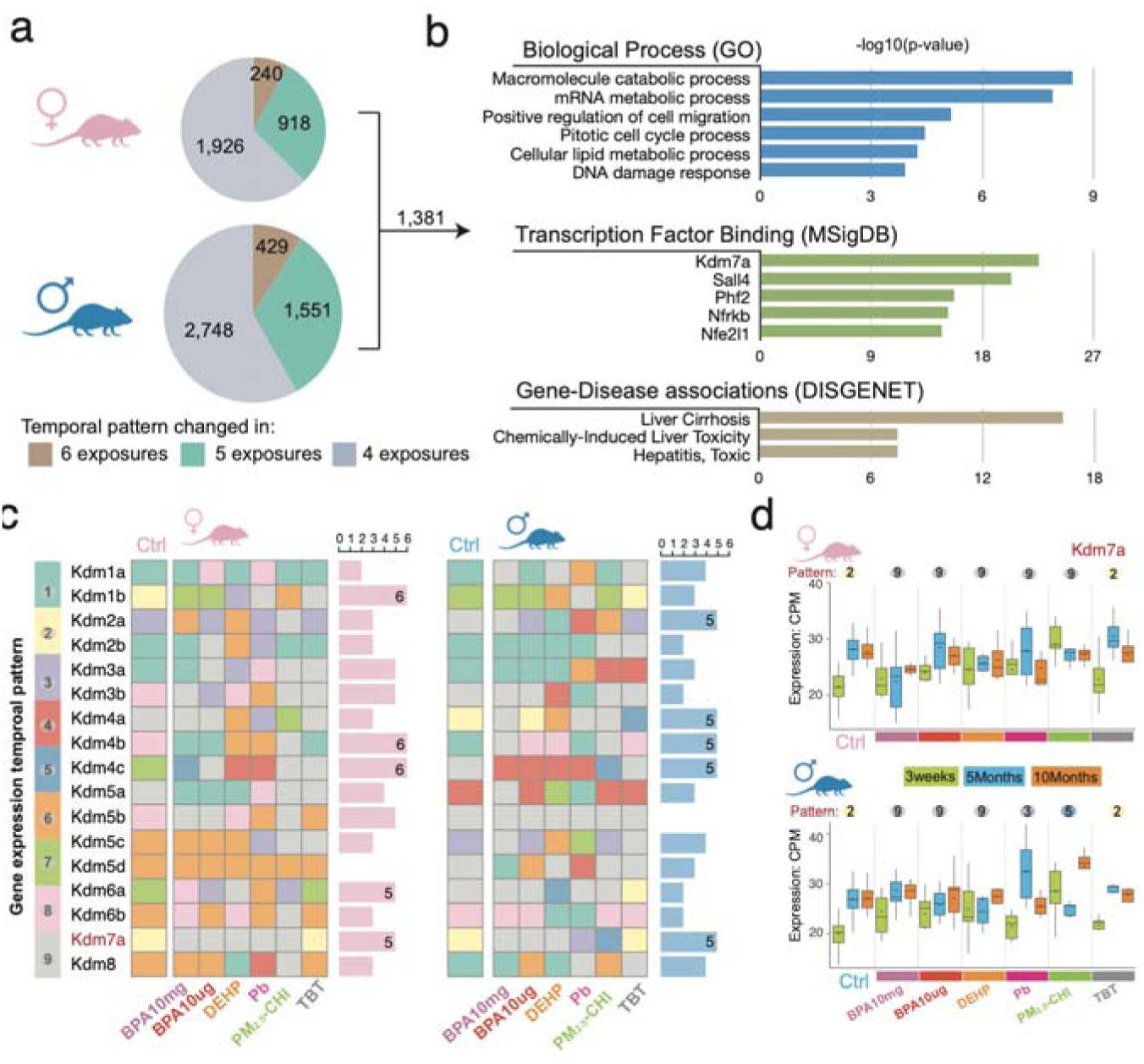
Common temporal changes in response to environmental exposures. **a)** genes with disrupted temporal expression patterns in at least 4 exposures; **b)** enriched biological processes and diseases of 1,381 commonly changed expression temporal patterns; **c)** temporal expression patterns of the *Kdm* family genes in each exposure; **d)** expression of *Kdm7a* in female (upper) and male (lower) liver.

### Toxicant-specific epigenomic and transcriptomic signatures

In the liver, we identified a total of 5,608 DEGs in female and 7,483 DEGs in male livers at weaning, young, or later adulthood that responded to at least one exposure [**S-Figure 6a**; **Figure 3a, 3b**]. These DEGs were distributed throughout the genome [**Figure 3c]**. Although the majority of DEGs were exposure-specific, 15-24% of DEGs at any of the three ages examined (3 weeks, 5 months, or 10 months) were targeted by multiple exposures, indicating both shared and distinct targets for these toxicants [**S-Figure 7**]. For example, 274 genes on chromosome 3were differentially expressed at 5 months in response to at least one exposure, and 58 of these responded to multiple exposures with distinct regulation patterns [**Figure 3d**]. As illustrated for *Car3*, which was downregulated in As-, BPA10µg-, Pb-, and TCDD-exposed adult male liver tissue [**Figure 3d, e**], decreased expression was accompanied by decreased chromatin accessibility in the region proximal to its transcription start site (TSS) [**Figure 3f**]. In adult female liver tissue, the *Gstm3* gene was down-regulated in response to As and PM_2.5_-JHU exposures but up-regulated in response to BPA10µg, PM_2.5_-CHI, and TBT [**Figure 3d**].

**Figure 6.**
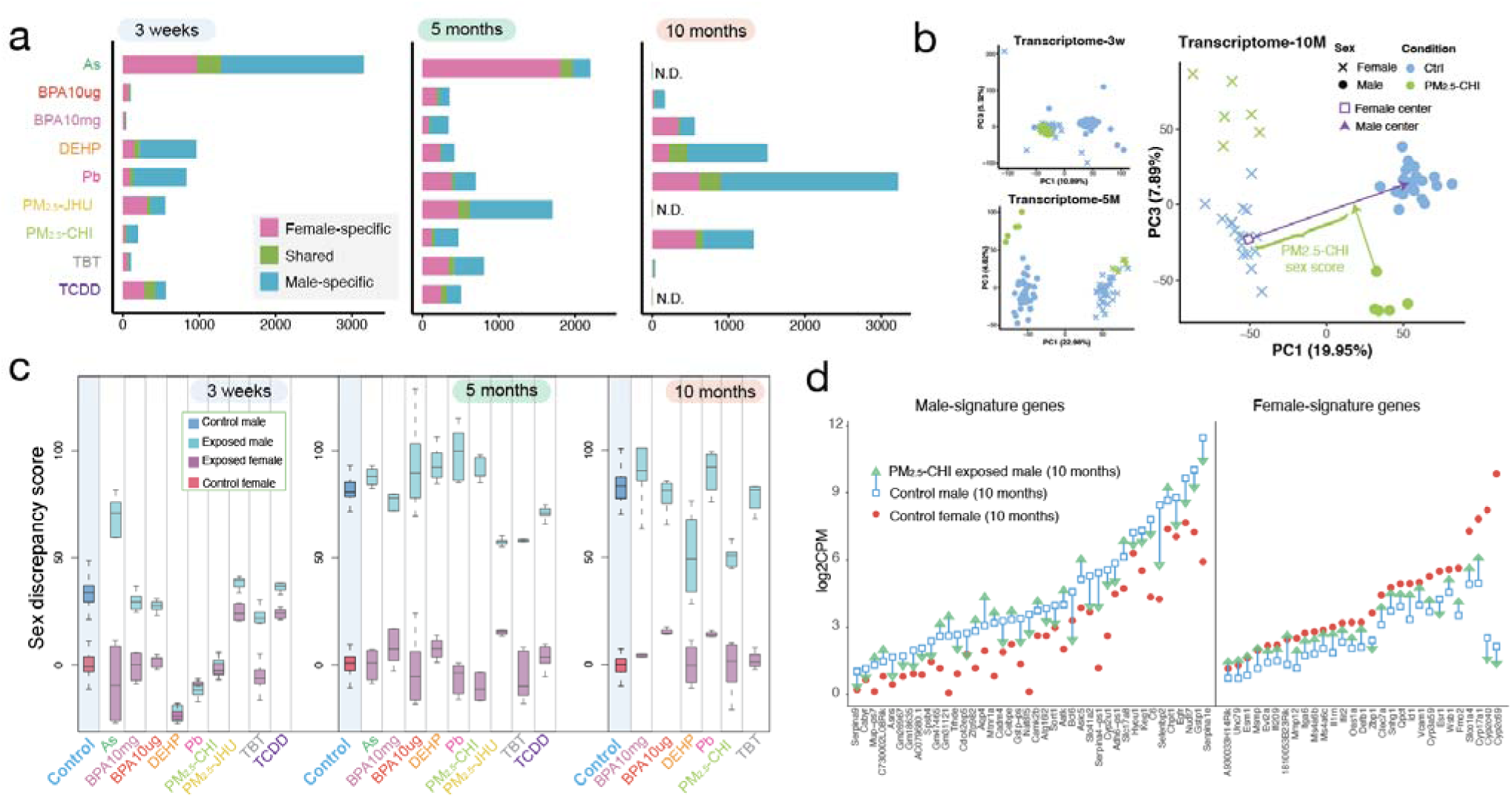
Sex-specific responses in mouse liver. **a)** female- and male-specific and shared responding transcriptomic signatures in both sexes to all exposures at three life stages; **b)** 2D PCA illustration of transcriptomes at 3 weeks, 5 months, and 10 months, with the demonstrated sex score measurement (demo) of one PM_2.5_-CHI-exposed male; **c)** Sex discrepancy scores of control and exposed mice at 3 weeks, 5 months, and 10 months; **d)** Expression changes of male-biased signatures (left) and female-biased signatures (right) in male livers at 10 months after early-life PM_2.5_-CHI exposure.

**Figure 7.**
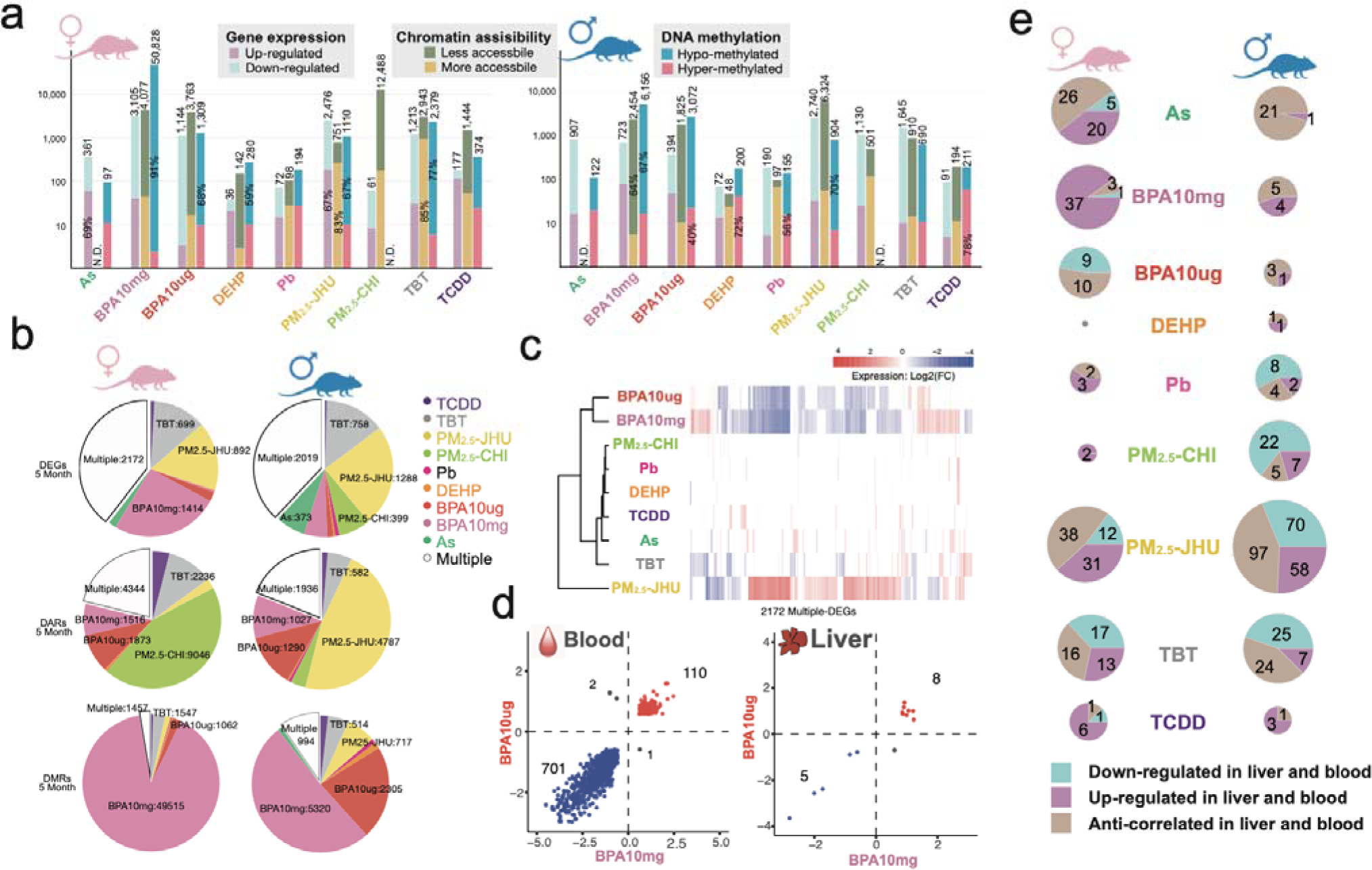
Molecular responses to environmental exposures in surrogate blood cells. **a)** number of differentially expressed genes, accessible regions, and DNA methylated regions in every exposure condition in female blood (left) and male blood (right); **b)** toxicant-specific and multiple-exposure-shared signatures of the transcriptome (top), chromatin accessibility (middle), and DNA methylation (bottom) in blood at 5 months; **c)** expression changes of multiple-exposure-shared genes of exposure signatures in 5 months female blood; **d)** expression changes of shared genes between BPA10mg signatures and BPA10µg signatures in female blood (left) and liver (right); **e)** numbers of shared DEGs between liver and blood in each exposure condition. Area size is normalized by log2(gene numbers).

Interestingly, differential gene expression at one age did not necessarily predict altered expression at other ages [**S-Figure 6a**]. Likewise, the number of DARs induced by toxicants did not necessarily predict the number of DEGs. For example, in As-exposed males, at weaning, 1,785 As-specific DEGs were identified, accompanied by robust increases in DARs. However, this response did not persist into young adulthood, where only 230 DEGs were identified at 5 months, accompanied by an equally dramatic decrease in chromatin accessibility [**Figure 3b, S-Table 3**]. Meanwhile, in As-exposed females, which exhibited a similar number of DEGs (1,201) at weaning, very few changes in chromatin accessibility were seen (450 DARs). At 5 months, the number of DEGs was relatively unchanged (1201 vs 1,467), but was maintained in the presence of a dramatic increase in DARs (from 450 to 10,071) [**Figure 3a]**. In DEHP-exposed males, few DEGs were identified at weaning (303 DEGs) or young adulthood (82 DEGs), but in later adulthood, 989 DEHP-specific signature DEGs were identified, although few DARs were identified at any age [**Figure 3a, S-Table 3].** This suggests other exposure-related changes, for example, altered transcription factor activity, could be driving changes in the transcriptome independently of changes in chromatin accessibility.

Notably, changes in chromatin accessibility in response to toxicant exposures were not necessarily directly correlated with changes in epigenetic modifications [**S-figure 7**]. For example, the gain or loss of open chromatin in exposure-induced DARs was not always accompanied by commensurate global changes in DNA methylation [**Figure 3a**, **b].** Furthermore, even in exposure settings where massive numbers of DARs were observed in response to toxicants, the majority of the genome retained the chromatin state of their matched vehicle controls [**S-figure 8**]. This was also the case when looking at specific DARs. In young adulthood, nearly half of the DARs in male livers exposed to BPA10mg (5,073 out of 10,444) exhibited changes in active chromatin marks for ChromHMM-defined chromatin states. In contrast, only ∼15% of PM_2.5_-CHI-induced DARs (685 out of 3,742) showed chromatin state changes commensurate with a switch to active chromatin [**S-figure 9; S-Table 4**].

Thus, parallel multi-omic profiling revealed that toxicant-induced changes in gene expression, chromatin accessibility, and DNA methylation did not always occur in lockstep [**Figure 3a,b**]. While the relationships between open chromatin and transcription, as well as DNA methylation and gene silencing, are well established, we observed many instances where toxicant-induced alterations in the transcriptome were not accompanied by commensurate changes in the epigenome, and vice versa. For example, in 5-month-old Pb-exposed female livers, 230 Pb-specific DEGs were identified, yet little change was observed in chromatin accessibility (57 DARs) or DNA methylation (193 DMRs). Conversely, significant alterations in the epigenome did not always correspond to substantial transcriptomic changes. For instance, in females exposed to BPA at 3 weeks (10 µg) and 5 months (10 mg), many DARs were identified (3,275 and 3,029, respectively), but minimal changes in gene expression occurred. Another striking example was seen in females exposed to TBT, where 32,875 differentially methylated regions (DMRs, including 29,983 TBT-specific) were observed at 5 months, yet many fewer signature DEGs (425 DEGs at 20 weeks [**S-Table 3**], including 136 TBT-specific DEGs) were identified in the livers of these animals [**Figure 3a, b**]. Thus, RNA-seq, ATAC-seq, and WGBS each yielded distinct insights into the persistent effects of early-life exposures on the transcriptome and epigenome.

### Early-life exposures reprogram the expression trajectory of signature genes

The longitudinal experimental design of TaRGET II presented a unique opportunity to explore temporal gene expression patterns and the impact of exposures on the expression trajectory of target genes. Our initial analysis of gene expression patterns as a function of age in vehicle controls identified nine reference temporal patterns for how gene expression changed between weaning, young, and later adulthood, identified as patterns 1 through 9 in **Figure 4a** [**S-Figure 10a, b**] (Methods). In terms of the overall trajectory of genes differentially expressed as a function of age, Patterns 1, 6, and 8 were defined by genes that decreased expression between weaning and later adulthood, while Patterns 2, 5, and 7 were defined by genes that increased expression over this interval.

To assign potential functionality to these patterns, overrepresentation gene ontology analysis (ORA) was performed to ask if genes within these 9 patterns represented biological functions and/or pathways [**Figure 4b**, **S-Figure 10c, S-Table 5**]. Remarkably, we found a significant association between many of these temporal expression patterns and specific biological functions. For example, in females, Pattern 1 genes, which are highly expressed at weaning, but then decline with age, were significantly associated with the cell cycle (p<10^-30^), as well as DNA replication, cytoskeletal organization, and cell adhesion. This decreased expression of genes involved in cell cycle and DNA replication is consistent with reduced cell proliferation as the liver ages, and diminished expression of cytoskeleton and cell adhesion genes associated with normal decreases in extracellular matrix production by nonparenchymal cells^66–68^. Pattern 2 genes, which increased between weaning and adulthood, were significantly associated with metabolism (p<10^-20^), catabolic processes, and interferon production, consistent with increased liver metabolism with age^69,70^.

To classify liver gene expression dynamics in response to early-life exposures, we next examined how exposure to toxicants affected the expression patterns of genes in Patterns 1 through 9. We observed significant disruption of normal temporal expression patterns due to BPA (10 mg and 10 µg), DEHP, Pb, PM_2.5_-CHI, and TBT [**Figure 4c**]. Early-life exposure to these toxicants disrupted between 25% and 90% of the target gene’s temporal expression patterns. This is illustrated for each exposure with pie charts showing the extent of gene disruption within each temporal expression pattern [Figure 4c, S-Table 6]. The degree of disruption varied depending on both the type of exposure and sex, but it was generally more extensive in males than in females. For instance, in males, DEHP disrupted the temporal expression of the majority of genes in Pattern 2 (83%), Pattern 4 (88%), and Pattern 5 (85%), while sparing more genes in Pattern 1 (41%) and Pattern 6 (47%) [**Figure 4c**]. This suggests potential impacts on liver metabolism and catabolism ^69,70^ (Pattern 2 genes), RNA translation^71^ (Pattern 4 genes), and energy metabolism^72^ (Pattern 5 genes) in these exposed males. In contrast, fewer genes within these same patterns were affected in females, who primarily showed changes in Expression of Pattern 2 (47%), Pattern 4 (60%), and Pattern 5 (46%) genes when exposed to DEHP. In females, BPA10mg, BPA10µg, Pb, and PM_2.5_-CHI had the most significant effects on the temporal gene expression patterns of genes in Patterns 2 (57%), 9 (56%), 4 (61%), and 5 (54%) [**Figure 4c, S-Table 6**].

The alluvial plots in **Figure 4d** provide a more detailed view of how the expression of genes in each of the nine temporal expression patterns changed as a function of exposure and age (3 weeks, 5, and 10 months), as illustrated in detail for BPA10µg, Pattern 9 genes in females and Pattern 1, 8 and 9 genes in males exposed to TBT. In females, Pattern 9 was defined by 2,536 genes whose expression normally did not change with age [**Figure 4a**]. However, in response to early-life exposure of BPA, 55% of these genes gained a different temporal expression pattern, now displaying the temporal expression of genes in Patterns 1, 3, 4, 7, or 8. In females exposed to BPA, 156 Pattern 9 genes, which normally would not change with age, progressively increased expression with age, thus acquiring a Pattern 7 (increase with age) trajectory [**Figure 4d**]. Remarkably, many of these reprogrammed Pattern 9 genes had similar functionalities– fatty acid oxidation and lipid metabolic processes – as the *bona fide* Pattern 7 genes [**S-Figure 10d**, compared to **Figure 4b**]. Other reprogrammed Pattern 9 genes that acquired Pattern 1 and 8 (decrease with age) trajectories, which were enriched for immune signaling, suggested the exposure-induced decline in immune function along with aging^73–75^ [**S-Figure 10d**].

In TBT-exposed males [**Figure 4e]**, 930 Pattern 9 genes acquired the temporal expression pattern of Pattern 2 (223 genes) and 3 (707 genes), becoming upregulated after weaning. These reprogrammed genes were enriched in cellular autophagy, lipid metabolism, and the TORC1 signaling pathway, which also characterized the *bona fide* Pattern 2 and 3 genes [**S-Figure 10c, d**]. A total of 600 Pattern 1 genes adopted the temporal expression profiles of Pattern 8 (373 genes), which were elevated in weanlings but declined with age, and Pattern 9 (227 genes), which exhibited a loss of age-related dynamic regulation. These Pattern 1 genes were enriched for apoptosis, DNA damage, and cytoskeleton remodeling functions, indicating that TBT exposure had a long-term impact on adult livers [**Figure 4e, S-Figure 10d**].

We extended this temporal pattern analysis further by examining sex-specificity in genes that were targeted by one or more exposures. As shown in **Figure 5a**, we found the temporal expression of 240 and 429 genes in females and males, respectively, was disrupted in response to all 6 early-life exposures; 3,084 and 4,728 genes were disrupted by at least 4 exposures in females and males, respectively. Of these, the temporal expression of 1,381 genes was disrupted in both males and females. Gene ontology analysis showed these genes were associated with mRNA and lipid metabolism, cell migration, cell cycle, and DNA damage response [**Figure 5b**], suggesting these pathways may be vulnerable targets in both sexes. Notably, these genes are significantly associated with liver diseases, including liver cirrhosis, toxic hepatitis, and chemical-induced liver damage^76–79^ [**Figure 5b**]. Interestingly, many were targets of specific transcription factors and chromatin modifiers, including histone demethylase *Kdm7a*, *Phf2, Sall3*, *Nfe2l1,* and *Nfrkb* [**Figure 5b**], which can actively respond to distinct environmental exposures^80–90^, suggesting that altered transcription factor activity and/or chromatin remodeling could be playing a role in how these environmental exposures were driving changes in gene expression.

The enrichment for *Kdm7a* target genes prompted us to investigate whether the temporal expressi n patterns of other chromatin modifiers were disrupted by early-life exposures, especially since these genes can remodel chromatin and amplify downstream effects on the expression of other genes. Indeed, the temporal expression of many histone demethylases in the Kdm family was altered by multiple early-life exposures [**Figure 5c**]. *Kdm1b*, *Kdm4b,* and *Kdm4c* showed changes in their temporal expression patterns in female livers in response to all six exposures. In male livers, the expression patterns over time for *Kdm2a*, *Kdm4a*, *Kdm4b,* Kdm*4c*, and *Kdm7a* were affected by five exposures [**Figure 5c**].

*Kdm7a* is a demethylase essential for tissue development that removes repressive H3K9me2 and H3K27me2 methyl marks, thereby facilitating the expression of previously silenced genes. Temporal expression *Kdm7a* normally follows Pattern 2 (i.e., increased expression with aging) in both female and male livers. In females, exposure to 5 toxicants shifted *Kdm7a* expression to a Pattern 9 pattern (i.e., no change in expression with age). However, the impact of this shift may have occurred much earlier, as for several exposures, the apparent lack of increase in adulthood was the result of a precocious elevation of *Kdm7a* expression in weanlings. In males, temporal expression of *Kdm7a* also normally follows Pattern 2, but in response to BPA and DEHP, it transitioned to Pattern 9, and in response to Pb and PM_2.5_-CHI, to Pattern 3 and Pattern 5, respectively [refer to temporal expression patterns identified in Figure 4 and expression levels by exposure and age in [**Figure 5d**]. These findings on *Kdm7a* highlight the importance of assessing gene expression as a function of age and illustrate both exposure-specific and sex-specific expression dynamics of genes targeted for reprogramming in the liver.

### Sex-specific epigenomic and transcriptomic reprogramming in response to environmental toxicants

The interesting sex-specific findings noted above, along with the power of TaRGET II data across multiple timepoints of exposure and both sexes, led us to examine in greater detail the similarities and differences in how males and females responded to these early-life exposures^91–98^. To do this, we first defined the exposure signature for each toxicant at weaning, early adulthood, and later adulthood, focusing on features shared by both sexes as well as those unique to each sex [**Figure 6a; S-Figure 6b, S-Figure 11**]. We observed significant sex-specificity, with only a small number of DEGs shared between females and males at any of the three ages. For example, As-exposure caused dysregulation of 1,283 genes in females at weaning and 2,207 in males, but less than 10% of these DEGs were common to both sexes. BPA10mg exposure induced almost entirely sex-specific signatures, with fewer than 2% of DEGs shared between males and females at any age [**Figure 6a**]. This pattern of pronounced sex-specific responses to early-life exposures was also reflected in DARs and DMRs [**S-Figures 11a, b**].

We further annotated changes in gene expression in males and females to determine if any specific pathways or biological processes were more or less prone to sex-specific reprogramming using GSEA [**S-Figure 12a**]. The immune-related pathways, TNFα and Interferon signaling, appeared to be especially vulnerable to reprogramming, being enriched in both sexes in response to multiple exposures. These pathways were negatively enriched in both female and male weanling liver transcriptomes in response to PM_2.5_-JHU, but positively enriched in later-adulthood in both sexes in response to TBT. They were also positively enriched in weanling female livers in response to both doses of BPA, but negatively enriched in later adulthood in female livers. DEHP exposure was associated with a reversed enrichment pattern of these immunity-related pathways between females and males. TBT exposure induced highly expressed genes enriched in the interferon-response pathways at all three ages in the female liver [**S-Figure 12a**]. Persistent disruption of immune signaling gene expression could potentially change the inflammatory milieu of the liver, although no phenotypic assessment of inflammation was conducted as part of TaRGET II studies. Importantly, however, the frequency with which these pathways were reprogrammed, albeit in different directions and in different sexes, suggests an inherent vulnerability of genes in these pathways targeted by multiple exposures.

To better understand how early-life exposures disrupted sex-specific exposure patterns, we defined a sex discrepancy score by measuring the differences in gene expression between male and female transcriptomes (with female as origin points in **Figure 6b**) at three ages shown in the PCA space (Methods). This difference is illustrated with the vector between female and male at 10 months in **Figure 6b**; an equivalent vector can be calculated between male and female at weaning and 5 months. We then calculated the sex-discrepancy score deviation caused by each exposure in males and females (Methods). The sex discrepancy scores not only measure the toxicant exposure-induced transcriptomic differences in gene expression between males and females at all three stages, as shown by the separation between male and female controls at all three ages, but also reveal that these differences are established before puberty and persist significantly into early and later adulthood[**Figure 6c**].

Several exposures significantly changed the sex discrepancy score at different ages, as illustrated for PM_2.5_-Chi in [**Figure 6b and S-Figure 12b**] and for all exposures as shown in [**Figure 6c]**. A decreased sex discrepancy score (e.g., PM_2.5_-Chi at 10 months) occurs when the exposed mice become more female-like, and an increased score occurs when exposed mice become more male-like. In males, for example, this could be due to decreased expression of male-biased genes and/or increased expression of female-biased genes^99–102^, resulting in a feminized expression pattern, and vice versa for females. In weanlings, Pb and PM_2.5_-CHI caused a substantial decrease in sex discrepancy scores in male mice, indicating their transcriptome became closer to the female-like expression pattern [**Figure 6c, S-Figure 12c]**. DEHP-exposed mice even indicated a strong female-biased expression pattern in both males and females at the weaning stage [**Figure 6c, S-Figure 1 d]**. Conversely, at weaning, sex discrepancy scores of PM_2.5_-JHU- and TCDD-exposed female mice increased, due to the acquisition of a more male-like expression pattern in female livers [**Figure 6c**].

In adulthood, the sex discrepancy scores between males and females increased significantly, primarily due to well-established sex-biased expression patterns after puberty [**Figure 6c].** Additionally, fewer exposures led to a deviation in sex discrepancy score compared to weaning, with PM_2.5_- and TCDD-exposed males showing a decline at 5 months, and DEHP- and PM_2.5_-CHI-exposed males showing a decline at 10 months [**Figure 6c**], indicating their more female-like transcriptome during aging. We also identified the most highly divergent sex-biased genes at all three ages in control female and male livers [**S-table 7**], and examined how these sex-biased genes were impacted by different early-life environmental exposures [**S-table 8**]. As an example, PM_2.5_-CHI-exposed males dramatically reduced sex discrepancy scores in later adulthood [**Figure 6c]**, meanwhile, 39 male-biased genes, which are normally highly expressed in males compared to females, were perturbed by early-life exposure, and 26 of them showed a significant decrease in expression, approaching that of females[**Figure 6d**], resulting nearly 2/3 of these male-biased genes acquired a more female-like expression pattern. In contrast, for female-biased genes, which were normally expressed at a lower level in males than females, 25 out of 29 of these genes significantly increased expression, again acquiring a more female-like level of expression in exposed male livers [**Figure 6d**]. Both acquisitions of female-like expression patterns in male livers decreased the sex discrepancy score in PM_2.5_-CHI males. Genes driving this decreased sex discrepancy score included *Bcl6*, a sex-dependent key regulator controlling male fat metabolism and immune response^103–105^, and *Egfr*, which primarily functions to regulate cell proliferation, survival, and regeneration in the liver^106–109^, all lost their male-biased high-level expression. Thus, early-life exposure to PM_2.5_-CHI caused an overall feminization of the male liver transcriptome in later adulthood.

### Early-life exposure-induced molecular changes in surrogate blood tissue

One of the primary objectives of the TaRGET II consortium was to assess surrogate tissues for their capacity to reflect toxicant exposures that relate to effects on target tissues, such as the liver. Such correlative signatures in surrogate tissues could then be used to develop biomarkers of exposure and/or health outcomes^110–112^. We found that early-life exposures caused significant molecular changes in adult blood cells, as summarized by the changes observed at 5 months in the young adult blood transcriptome, DNA methylome, and chromatin accessibility in response to all nine exposures [**Figure 7a**]; detailed data are provided in S-Table 8. Most notably, we observed distinct patterns in how different “omic” layers of regulation responded to various exposures, including significant sex-specific differences and a frequent lack of agreement between changes detected across different omic platforms, as discussed previously for the liver.

For example, in the blood of females exposed to 10 mg of BPA, extensive DNA hypomethylation was observed (91% of the 50,828 differentially methylated regions compared to control animals). However, in the transcriptome, increases and decreases in gene expression were roughly equal, as were increases and decreases in chromatin accessibility **[Figure 7a].** These numerous changes across different “omic” layers indicate a systemic response to BPA exposure, including the lasting reprogramming of the epigenome and transcriptome in blood, and the potential to develop surrogate biomarkers. Importantly, while DNA methylation changes caused by early-life exposures were mainly unidirectional, changes in chromatin accessibility and gene expression were bidirectional. Therefore, this strong tendency toward hypomethylation was not accompanied by a corresponding shift in the transcriptome or chromatin accessibility, again emphasizing that each “omic” layer provides distinct molecular insights. Global changes in chromatin accessibility and the DNA methylome did not necessarily predict the transcriptional response to early-life environmental exposures. Meanwhile, persistent epigenetic changes observed in the livers of young adults following various exposures differed significantly from those in the blood. This difference is evident when comparing the data in **Figure 3a** from the liver to the blood data **in Figure 7a**. While many exposures caused notable changes in DNA methylation in blood—including strong hypomethylation in females exposed to BPA—only TBT induced a similar substantial change in the liver. Very little hypomethylation was observed in the female liver in response to BPA. In males, significant hypomethylation was detected in blood at 5 months in animals exposed to PM_2.5_-JHU and TBT, but few significant change was observed in the livers of these animals. Compared to the liver, DEHP and Pb exposures resulted in only minimal molecular changes in the blood tissue that serves as a surrogate.

Importantly, although not always consistent, persistent changes in the blood of one or more of these omic layers proved useful as a correlative signature of exposure, with many changes observed in one or more of these molecular readouts in response to early-life environmental exposures [**Figure 7b, S-Table 8**]. Additionally, as seen in the liver, these alterations in blood revealed distinct exposure-specific molecular fingerprints for various exposures that persisted into young adulthood, with a notable sex-specific pattern. For example, BPA exposure, especially at high doses, significantly altered the blood transcriptome and DNA methylome in both sexes, but with a stronger response in females, creating a unique “fingerprint” associated with this exposure. PM_2.5_-JHU exposure induced numerous DEGs and DARs in male blood, while PM_2.5_-CHI exposure caused a significant increase in DARs in female blood but not in males [**Figure-7b, S-table 8**]. These findings highlight the exposure- and sex-specific differences in persistent epigenetic and transcriptomic changes caused by early-life environmental insults.

Further exploration of blood signature DEGs revealed some surprising patterns [**Figure 7c, S-Figure 13a**]. High- and low-dose BPA exposure signatures in female blood shared many genes, which were also reprogrammed by PM_2.5_-JHU but in the opposite direction [**Figure 7c**]. This suggests that these genes may be particularly susceptible to reprogramming, although the effects on gene expression are exposure-specific. The strong similarity between transcriptomic changes caused by high- and low-dose BPA exposure resulted in 110 genes being co-upregulated and 701 genes being co-downregulated, a pattern not seen in female liver exposed to BPA [**Figure 7d]**. However, this co-regulation pattern induced by different BPA doses was not conserved in male blood. TBT had a closer relationship to PM_2.5_-JHU exposure, with 411 co-downregulated and 84 co-upregulated genes [**S-Figure 13b**].

Looking across all 9 exposures, we also assessed the overlap between blood and liver DEGs, identifying genes that were commonly increased, decreased, or showed anticorrelated expression patterns in the two tissues in young adults at 5 months [**Figure 7e, S-Figure 13c**]. We found that the number of overlapping genes, which were differentially expressed in both liver and blood, varied widely across exposures. PM_2.5_-JHU showed the greatest overlap between target and surrogate tissues in both males and females, while DEHP showed the least. In females exposed to BPA10µg, 94 liver DEGs were identified at 5 months, of which 41 (∼40%) were also differentially expressed in blood, and 38 of these (>90%) changed in the same direction in both tissues. Conversely, only 9 out of 256 male BPA10mg liver DEGs were also differentially expressed in blood at 5 months. In response to TBT, 46 out of 425 female and 56 out of 467 male liver DEGs were detectable in the blood of young adult mice. In DEHP-exposed animals, none of the 252 female liver signature DEGs and only 2 out of 188 male liver signature DEGs were also differentially expressed in the blood. These findings highlight the complexity of comparing cross-tissue transcriptomic signatures, even under well-controlled systemic exposure conditions and despite genomic homogeneity, as seen in the C57BL/6 inbred mouse strain used for these analyses.

## Discussion

Persistent changes in DNA methylation, histone modifications, and chromatin accessibility caused by early-life environmental exposures can influence gene expression patterns in adulthood and contribute to the development of diseases and long-term health issues^1,5,6^. The TaRGET II Consortium carried out the first multi-exposure, multi-omic longitudinal profiling studies to systematically investigate how early-life environmental exposures impact epigenetic modifications and transcriptomic dynamics later in life. The well-controlled exposure and profiling studies conducted by the Consortium have generated new high-resolution epigenomic maps and transcriptomic profiles of target and surrogate tissues from 3-week, 5-month, and 10-month-old mice. These datasets enable comparisons across different exposures, sexes, ages, and profiling methods. The standardized study design, which utilized a diverse range of environmental toxicants, including As, BPA, DEHP, Pb, PM_2.5_, TBT, and TCDD, offered novel and otherwise inaccessible insights into the molecular signatures induced by various exposures at different ages, tissues, and between sexes.

In the first target tissue examined under the TaRGET II project, the mouse liver, we discovered that each environmental exposure had a unique signature. Reprogrammed genes forming exposure signatures from different toxicants showed very limited overlap. This detailed profiling at the individual mouse level revealed that even for a single toxicant, there were age-specific and sex-specific differences in molecular signatures at both transcriptomic and epigenetic levels. Importantly, since all exposures occurred from pre-conception to weaning, and exposure ceased at the 3-week profiling point, disruption of the transcriptome and epigenome persisted into adulthood. This suggests a lasting reprogramming of epigenetic memory that extends beyond the immediate transcriptomic response to these environmental toxicants^113–115^. The data identified several pathways vulnerable to effects from multiple exposures: immune-related signaling^75,116,117^, liver metabolic processes^3,49,118^, transcription factor binding, especially chromatin remodelers such as demethylases^21,81,84,119^, and processes involved in liver diseases such as cirrhosis^120–122^.

Surprisingly, we found that data generated by different profiling approaches for various “omics”, including transcriptomic, epigenomic, and chromatin accessibility, did not provide redundant information. Instead, each “omic” layer provided unique data and insights, and together, they created a more comprehensive view of how early-life exposures affect the adult liver than any single assay alone. These data showed that the transcriptome’s response to early-life exposures could not always be predicted solely based on changes in the epigenome, and vice versa. In cases where persistent epigenomic reprogramming occurs without corresponding changes in the transcriptome, this reflects what has been described as “silent reprogramming” ^123^, which, although not immediately linked to changes in gene expression, may “prime” genes for abnormal expression in response to other environmental challenges later in life. When changes in gene expression are observed without concurrent changes in the epigenome, other factors such as the gain or loss of transcription factor activity at specific binding sites could be key drivers of those changes.

The longitudinal experimental design of TaRGET II, which involved collecting data from both males and females exposed equally (often within the same litter), allowed us to examine how environmental toxicants influence liver gene expression dynamics in the two sexes over time. We observed very clear, robust sex-specific differences in how the liver responded to these early-life exposures, even before puberty. These sex-specific responses affected different functional pathways in males and females, or sometimes the same pathway but in opposite directions depending on the. For instance, GSEA results showed that DEHP strongly activated numerous immune response and metabolic pathways in the livers of males in later adulthood, while the same pathways were repressed in females [**S-fig 12a**].

We also examined the commonality and differences between target tissues and surrogate tissues in response to toxic environmental exposures. We found significant molecular changes in blood that were unique to each exposure. These molecular fingerprints could serve as potential surrogate tissue biomarkers of exposure, and possibly response, for future research. We observed strong tissue-specific molecular responses to early-life exposure, with very little overlap between molecular changes in target and surrogate tissues. The toxicant-specific signatures identified in blood could enable the development of biomarker-based monitoring systems, even long after the exposure has occurred.

In summary, the TaRGET II consortium has developed a comprehensive multi-omics resource across multiple tissues at three life stages, following early-life exposure to nine distinct environmental toxicants in mice. This dataset provides invaluable benchmarking of transcriptomic and epigenetic profiles, which will support and enhance future environmental health research related to toxicant exposures.

## Acknowledgements

This work was supported by the NIEHS as part of the Toxicant Exposures and Responses by Genomic and Epigenomic Regulators of Transcription II (TaRGET II) Consortium through U24ES026699 (T.W.), U01ES026697 (D.D), U01ES026719 (C.L.W., M.S.B., and T.W.), U01ES02672 (S.B., S.R., and W.T.), U01ES026718 (G.M.), and U01ES026717(D.A.). This work was also supported by NIH through R35GM142917 (B.Z.), P30ES017885 and R35ES031686 (L.S, D.D.), R01 S028802 (J.C.), P30ES030285 (C.L.W.), U24HG012070 (T.W.), U41HG010972 (T.W.), and U24NS132103 (T.W.). This work was also supported by Chan Zuckerberg Initiative (B.Z.) and Diana Helis Henry Medical Research Foundation (69760-I, C.L.W.).

We acknowledge program leadership by members of the NIEHS TaRGET II workgroups, especially Fred. L. Tyson, Kim McAllister, Christopher G. Duncan, Amanda Garton, Lisa H. Chadwick, Maya Evanitsky.

## Competing interests

The authors declare no competing financial interests.

## Additional information

All the datasets, including both raw data and processed data, are available at TaRGET II data portal https://dcc.targetepigenomics.org/, and GEO accessions GSE146508.

All the analysis results are visualized at the accompanying database ToxiTaRGET https://toxitarget.com/.

## TaRGET II consortium(alphabetical)

### Integrative analysis leads and coordination

Benpeng Miao, Ting Wang, Bo A. Zhang,

### Integrative analysis

Cristian Coarfa, Justin A. Colacino, Shuhua Fu, Ravindra Kumar, Prashant Kumar Kuntala, Bongsoo Park, Wanqing Shao, Laurie K. Svoboda

### Integrative data production and processing

Gregory E. Crawford, Robert B. Hamanaka, Claudia Lalancette, Daofeng Li, Shaopeng Liu, Benpeng Miao, Heather B. Patisaul, Maureen A. Sartor, Tim Wiltshire, Xiaoyun Xing, Bo A. Zhang

### TaRGET II Data Production and Processing contributor

Nicole E. Allard, Raymond G. Cavalcante, Rengul Cetin-Atalay, Yujie Chen, Youngshim Choi, Alan Du, Elisa Ruiz-Echartea, Jackson P. Fredenburg, Tianyi Fu, Jaclyn M. Goodrich, Sandra L. Grimm, Silas Hsu, Brian M. Horman, Yiran Hou, Rahul Jangid, Yan Jin, Tamara R. Jones, Tiffany A. Katz, SunHong Kim, Prashant Kumar Kuntala, Yemin Lan, Bethany Latham, Tandao Li, Yan Li, Sydney Lierz, Siyu Liu, Juheon Maeng, Angelo Y. Meliton, Rachel K. Morgan, Kari Neier, Jackson P. Parker, Bambarendage P.U. Perera, Jayant M. Pinto, Deepak Purushotham, Dhivyaa Rajasundaram, Palanivel Rengasamy, Christine A. Rygiel, Alexias Safi, Erica Pehrsson, Kaitlyn A. Sun, Vinesh Vinayachandran, Kai Wang, Parker S. Woods, Bonnie HY Yeung, Jinhu Yin, Yu Zhang, Xiaoyu Zhuo

### Co-principal investigators

Cristian Coarfa, Gregory E. Crawford, Heather Lawson, Michael Province

### Scientific program management

Lisa H. Chadwick, Christopher G. Duncan, Kimberly A. McAllister, Frederick L. Tyson

### Principal Investigators

David Aylor, Marisa S. Bartolomei, Shyam Biswal, Dana C. Dolinoy, Gökhan M. Mutlu, Sanjay Rajagopalan, Wan-Yee Tang, Cheryl Lyn Walker, Ting Wang

## Notes

### Competing Interest Statement

The authors have declared no competing interest.

