## Supplementary material for "Comprehensive Transcriptomic and Epigenomic Insights into Environmental Toxicant Exposures: The TaRGET II Resource": S-Figure-1

Supplementary Figure 1

a

PCA plot for RNA-seq samples of 5 month liver

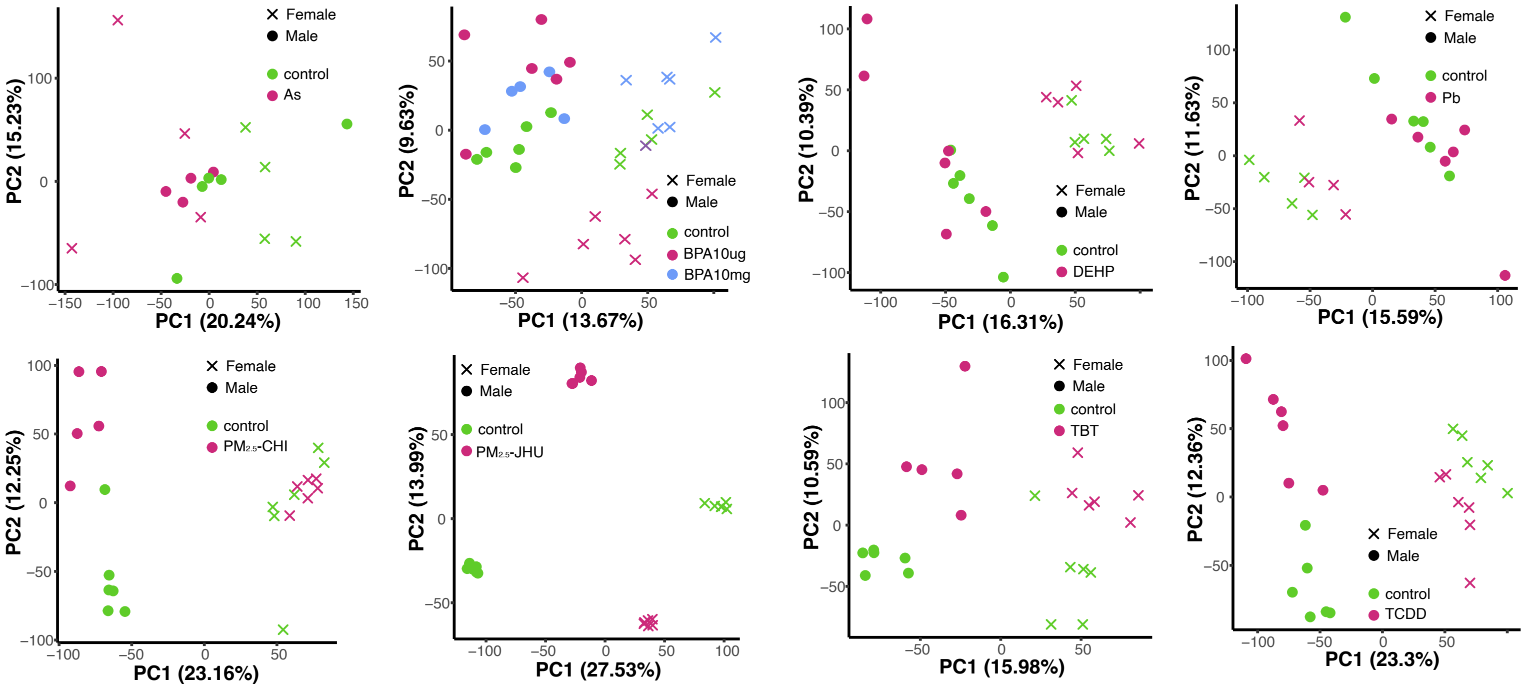


b

PCA plot for ATAC-seq samples of 5 month liver

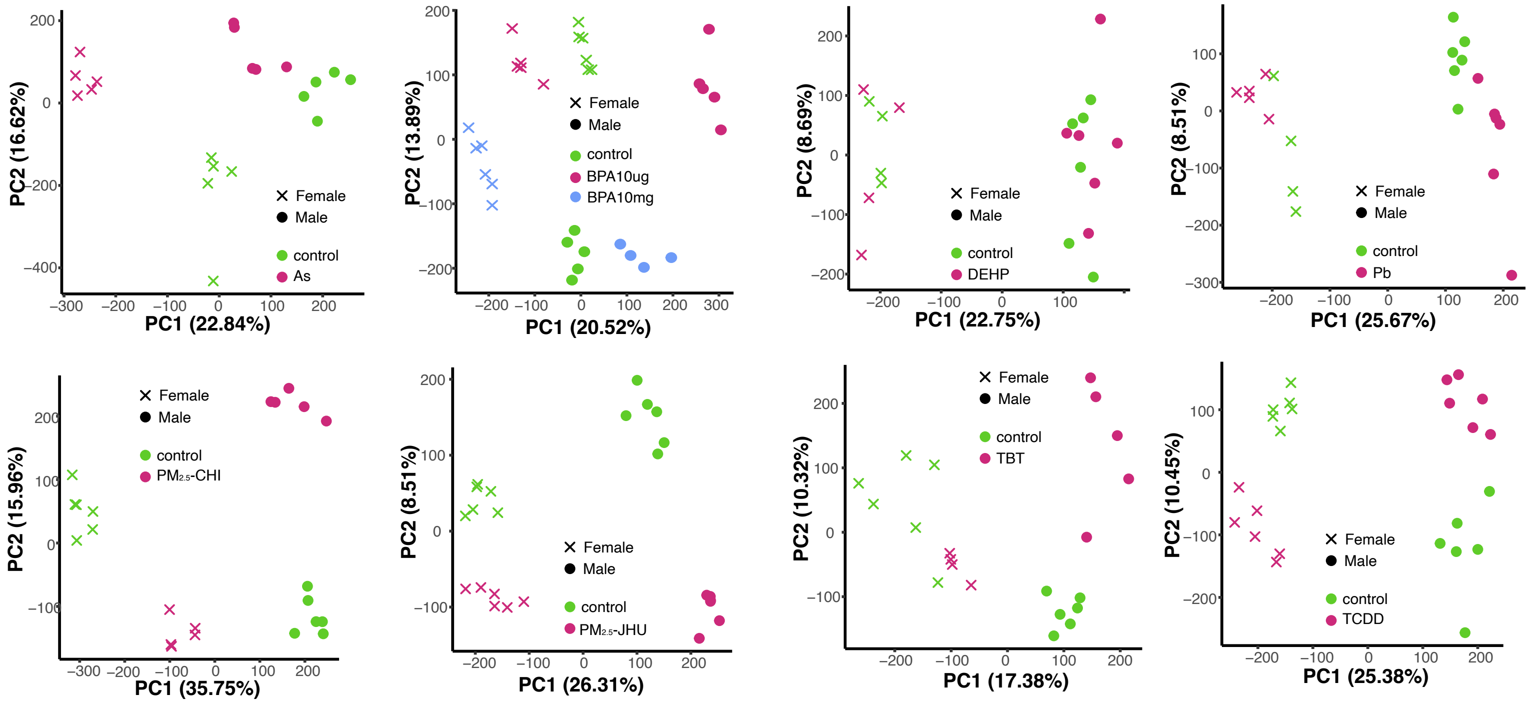
