## Supplementary figures and images for "Comprehensive Transcriptomic and Epigenomic Insights into Environmental Toxicant Exposures: The TaRGET II Resource"

### S-Figure-2

Supplementary Figure 2

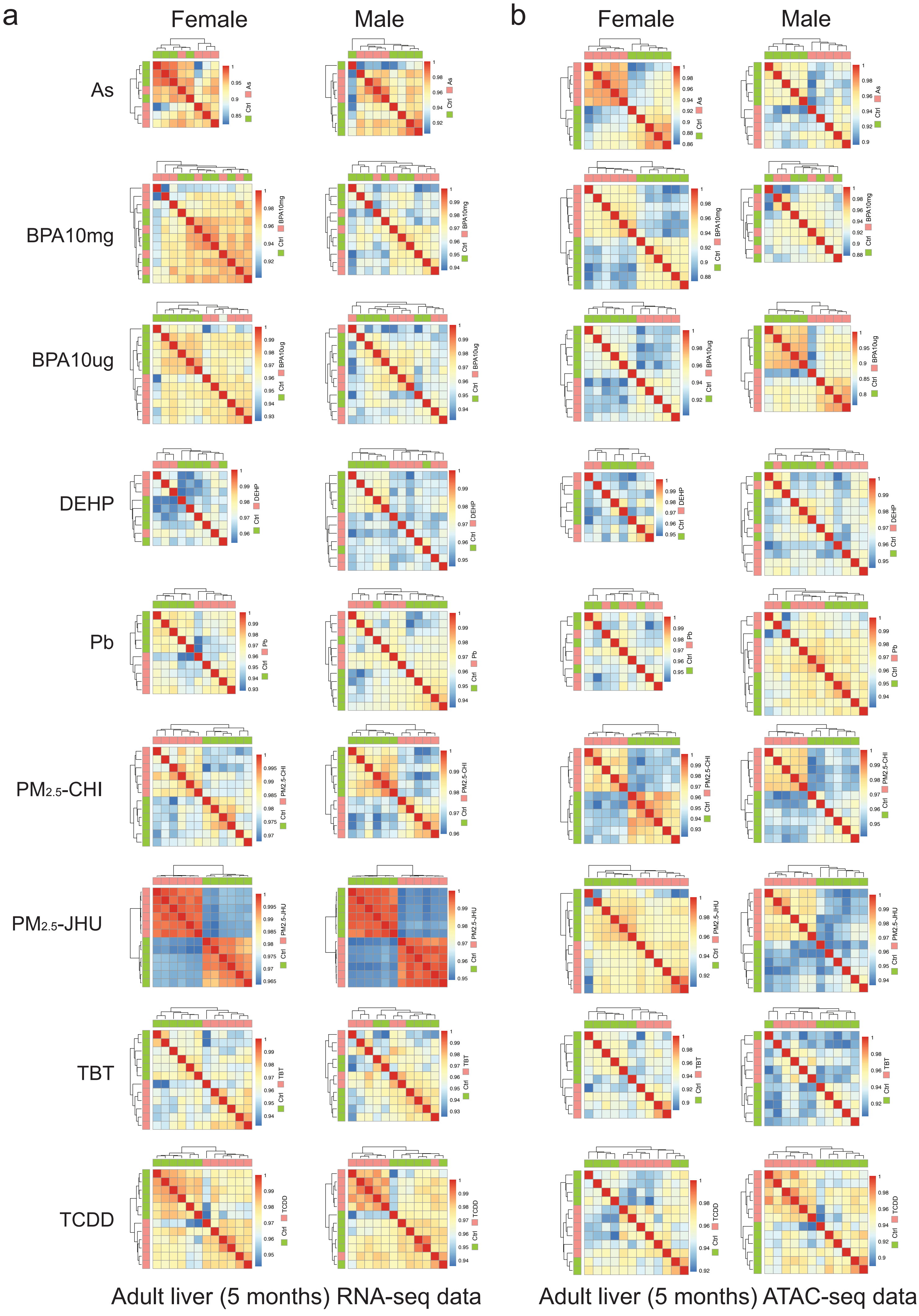

### S-Figure-4

Supplementary Figure 4

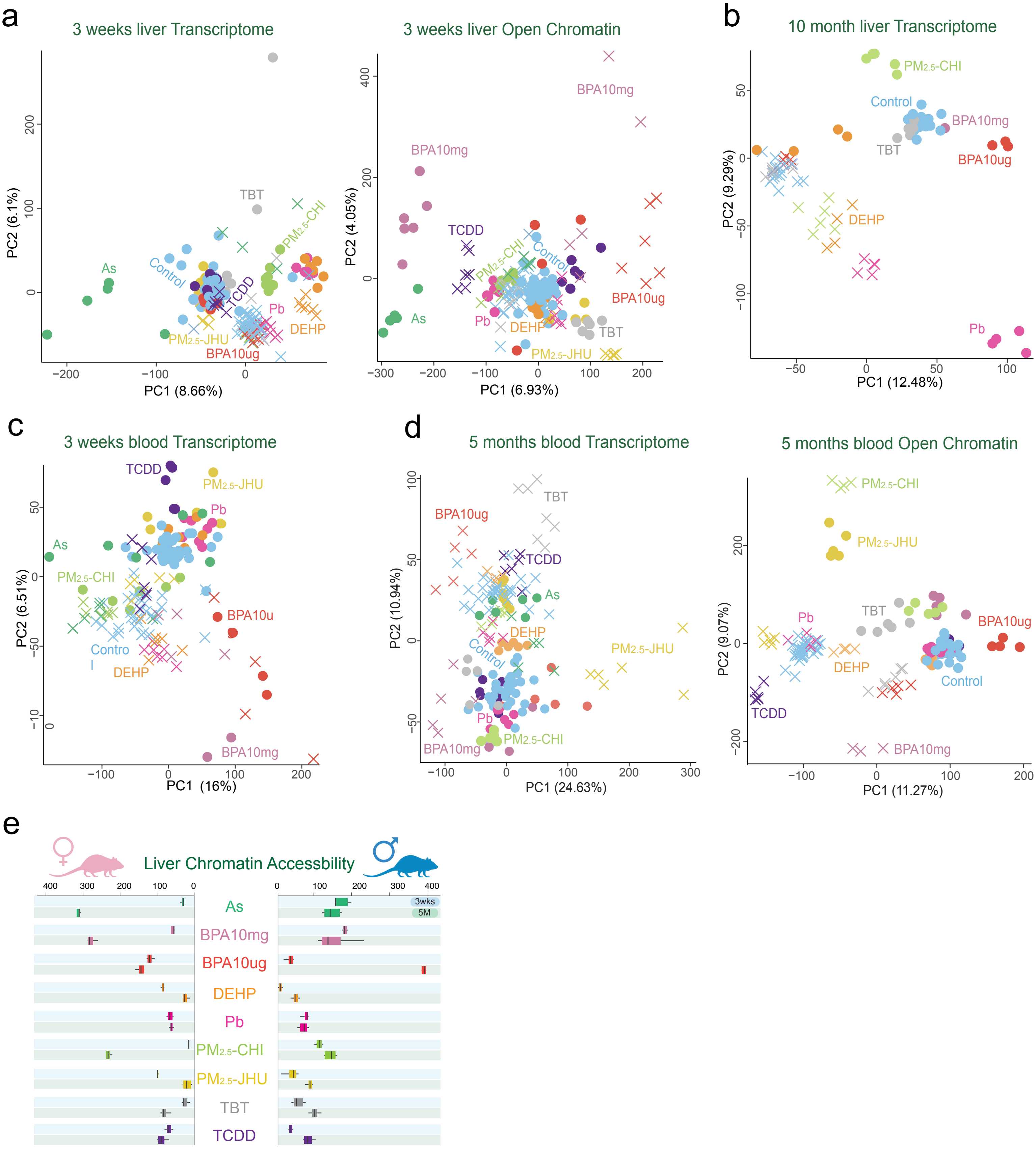

### S-Figure-5

Supplementary Figure 5

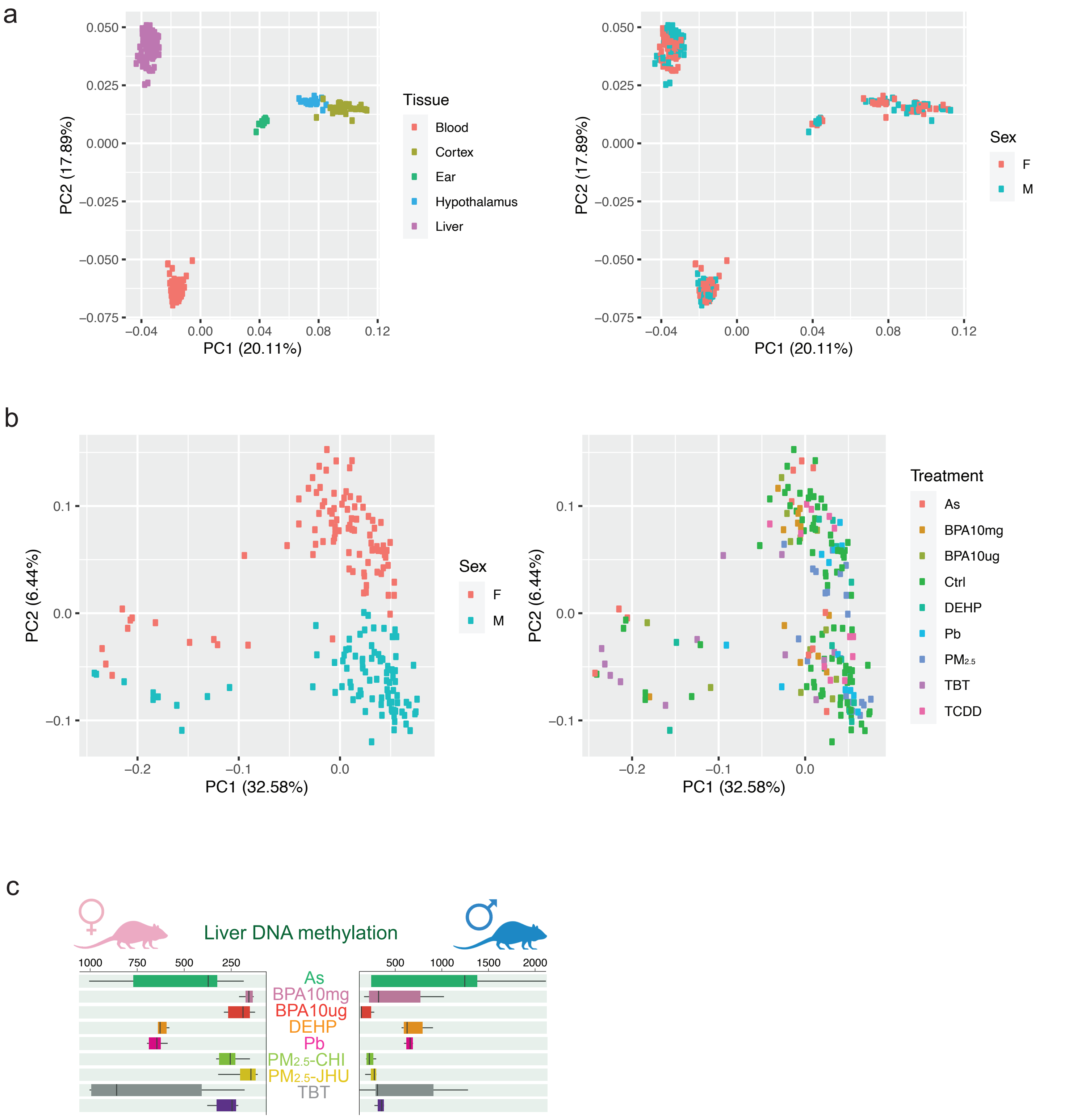

### S-Figure-6

Supplementary Figure 6

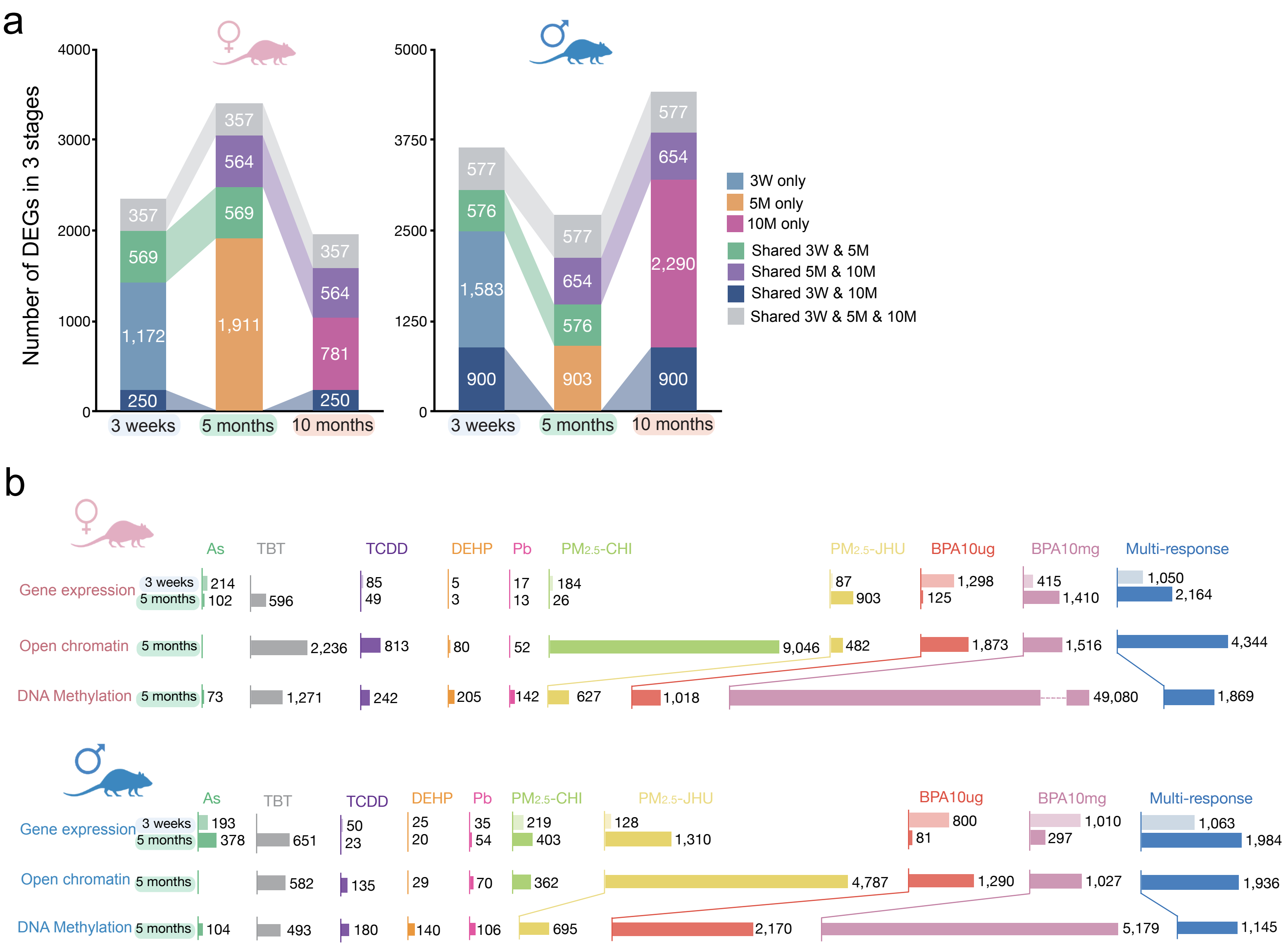

### S-Figure-7

#### Supplementary Figure 7

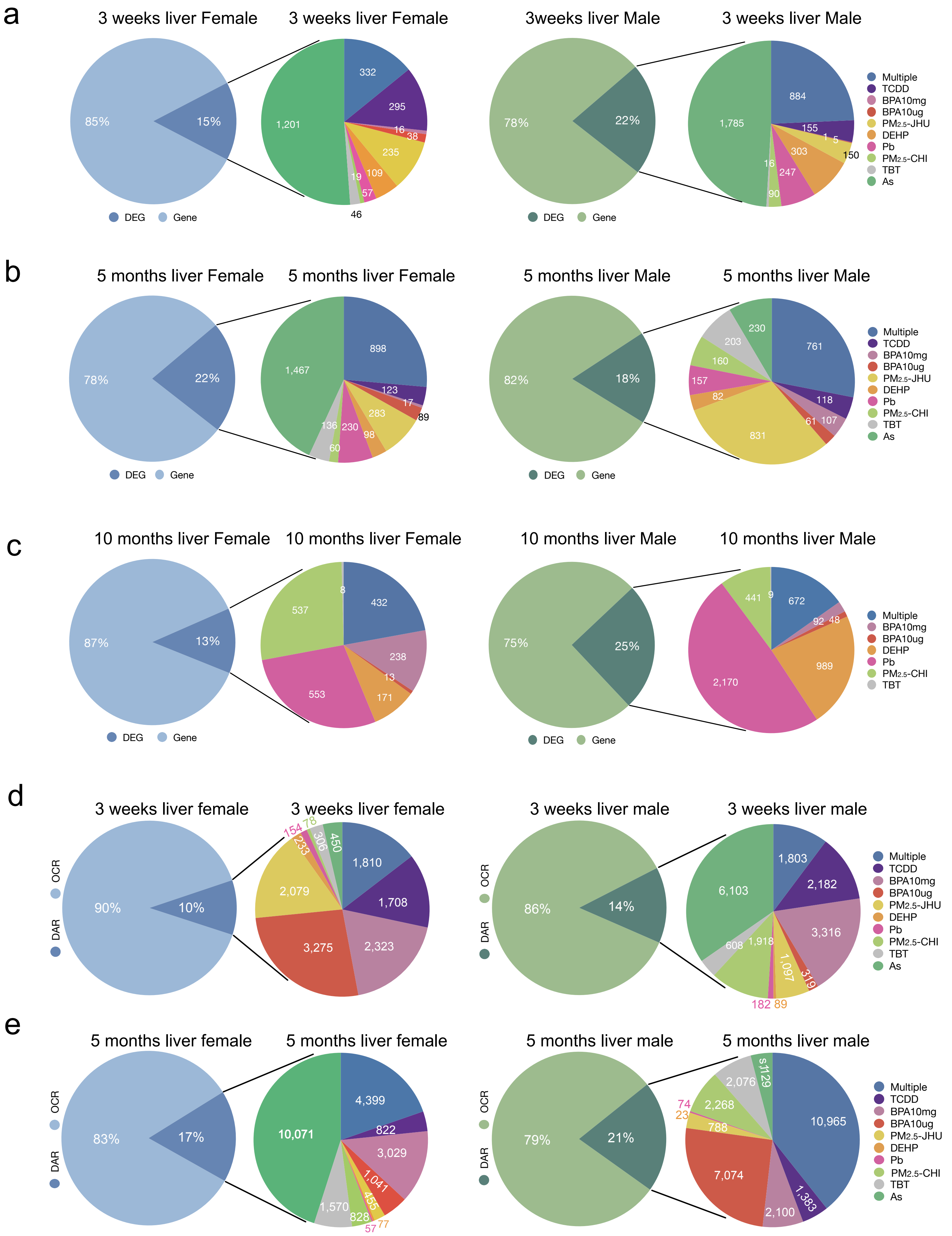

### S-Figure-8

Supplementary Figure 8

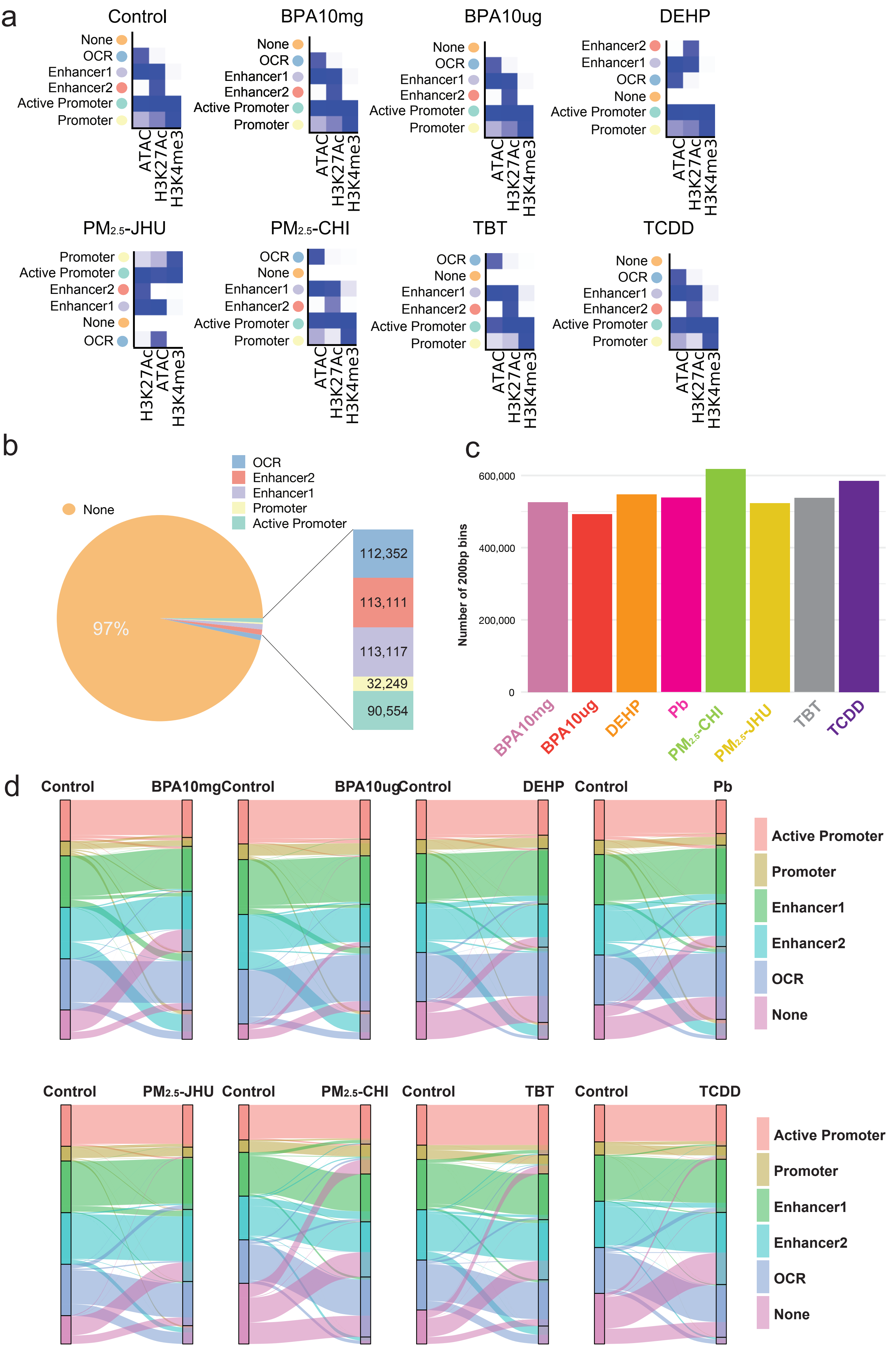

### S-Figure-9

#### Supplementary Figure 9

a

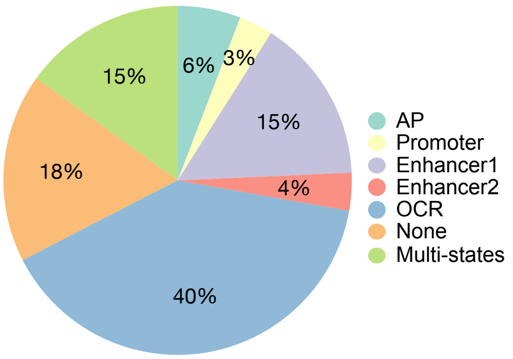


b

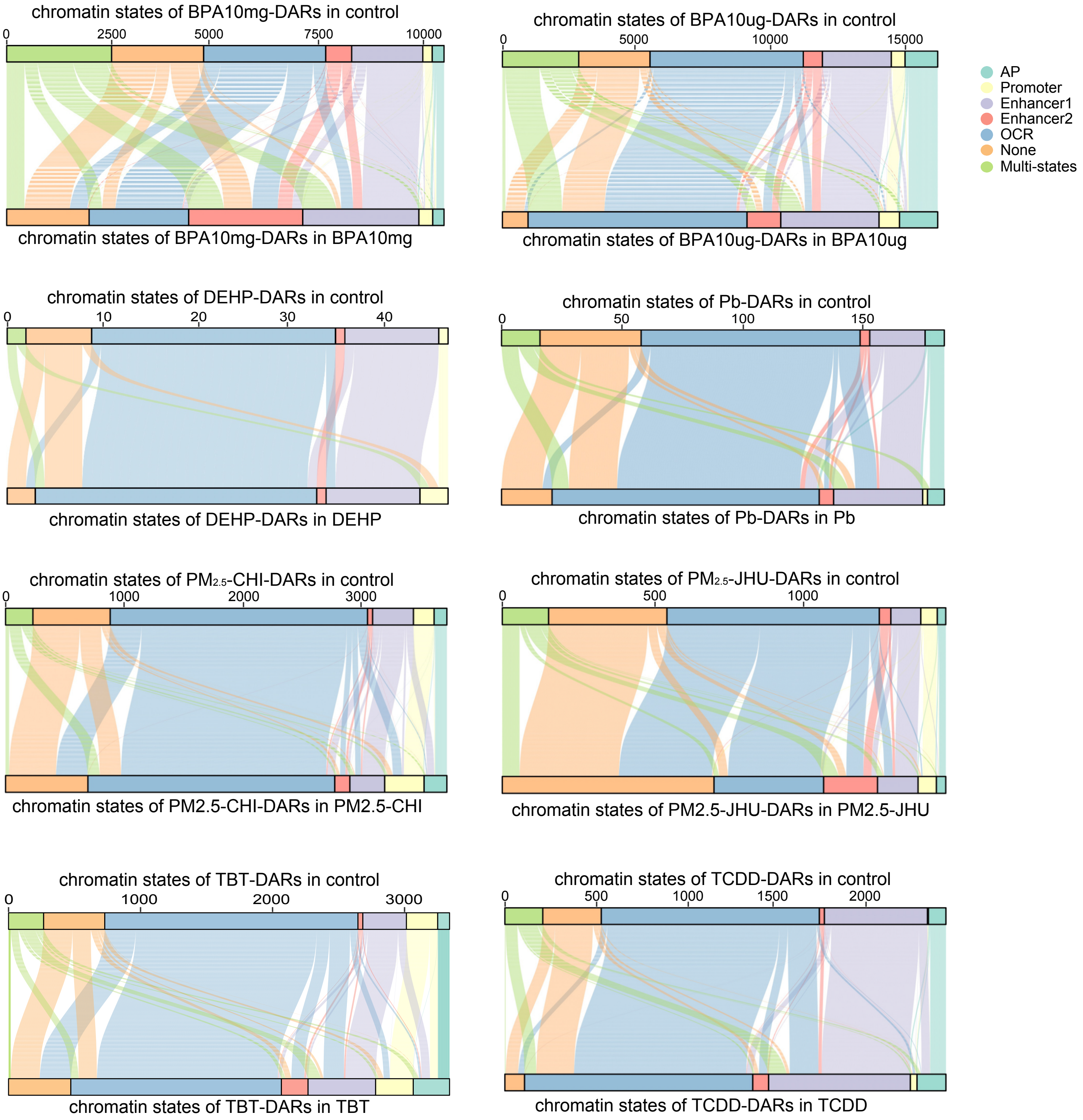

### S-Figure-10

#### Supplementary Figure 10

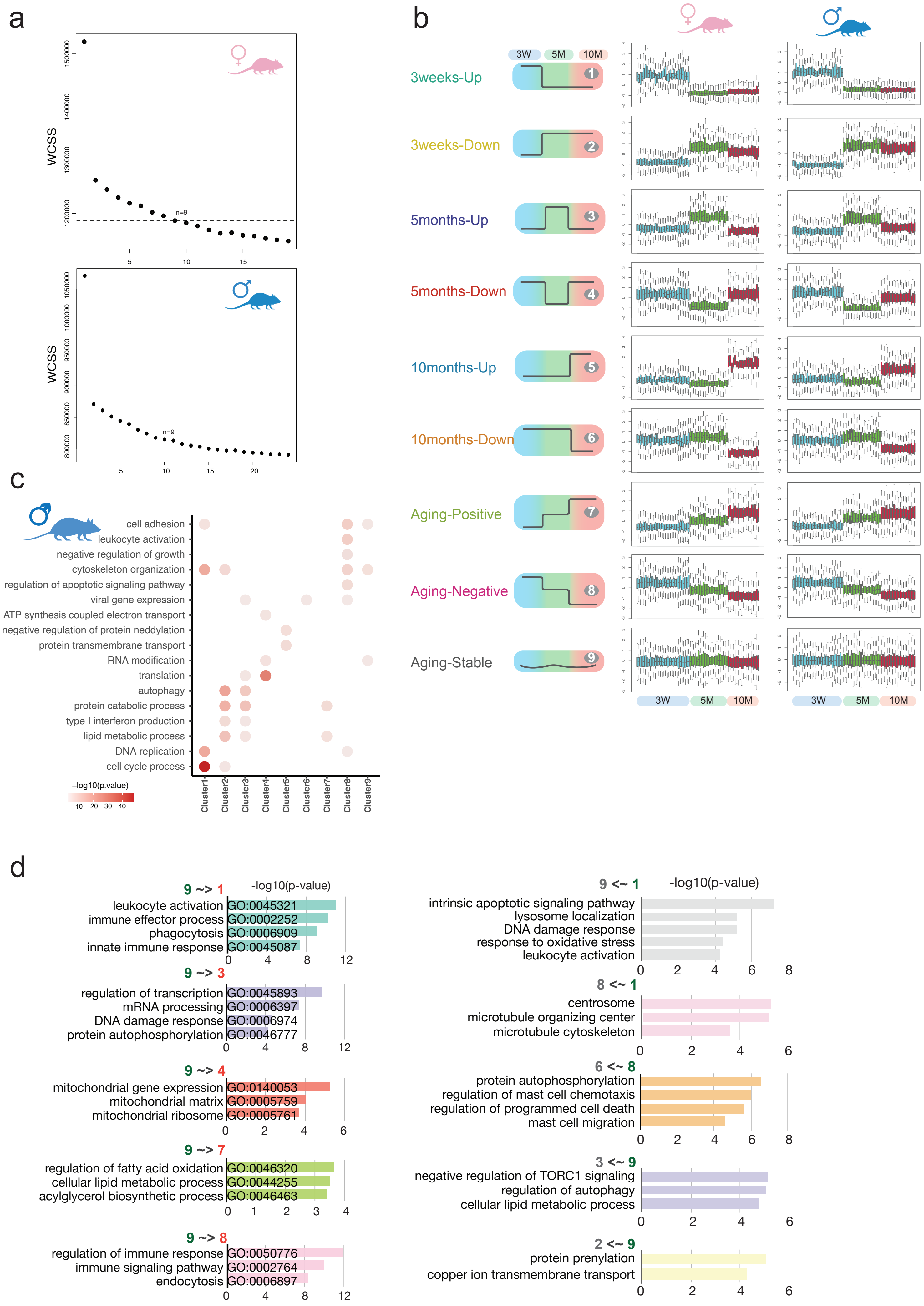

### S-Figure-11

##### Supplementary Figure 11

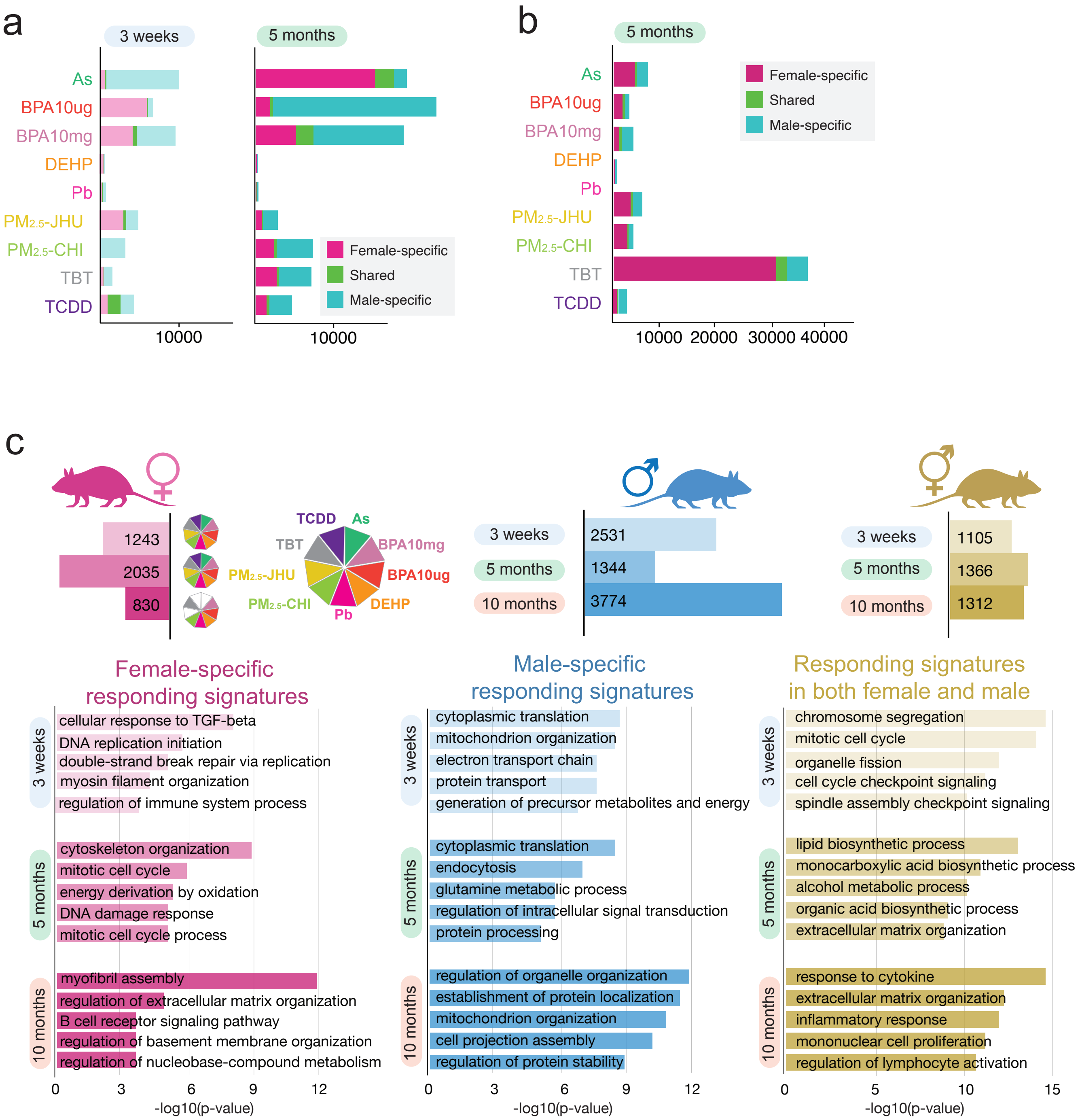

### S-Figure-12

#### Supplementary Figure 12

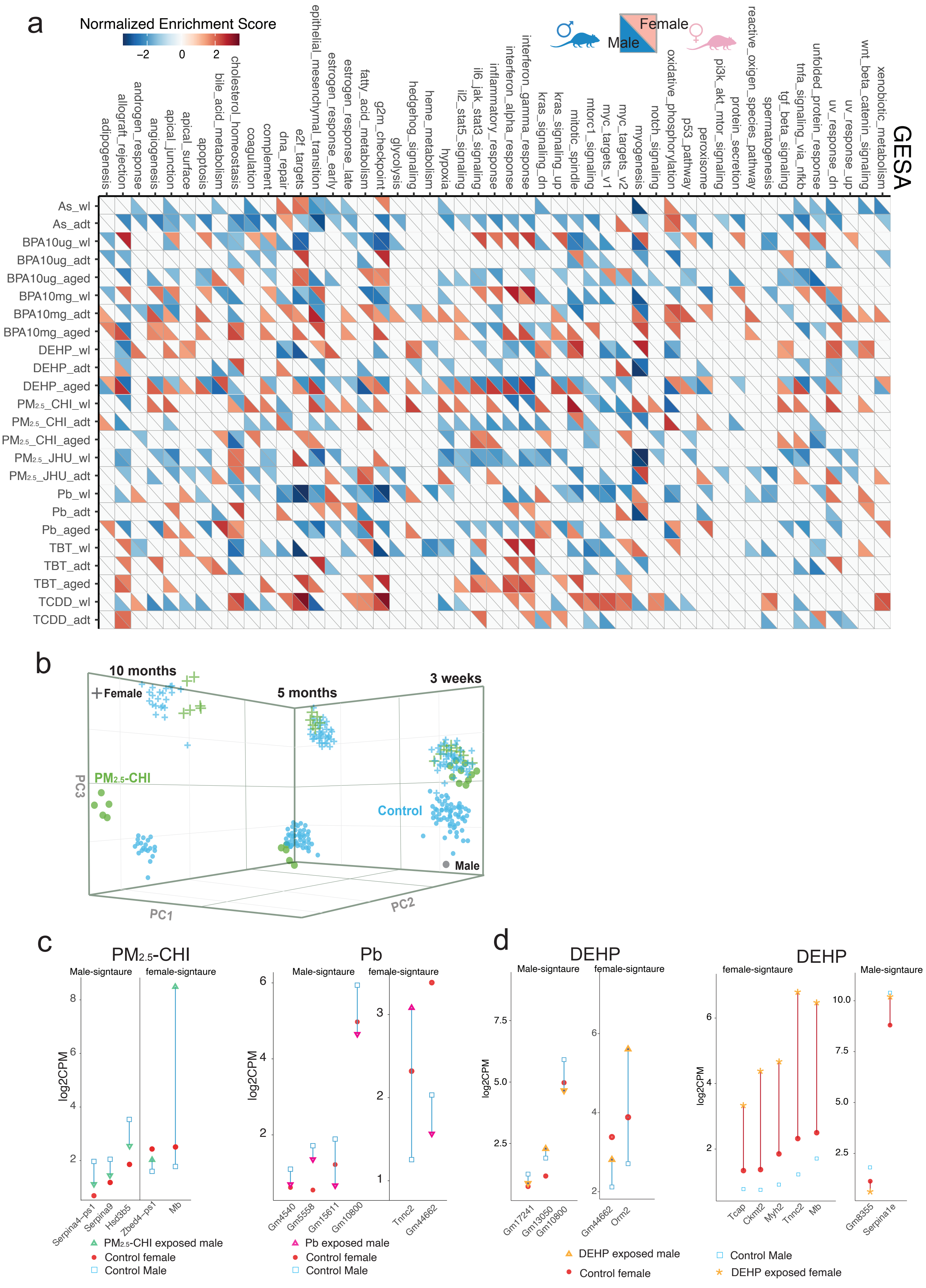

### S-Figure-13

#### Supplementary Figure 13

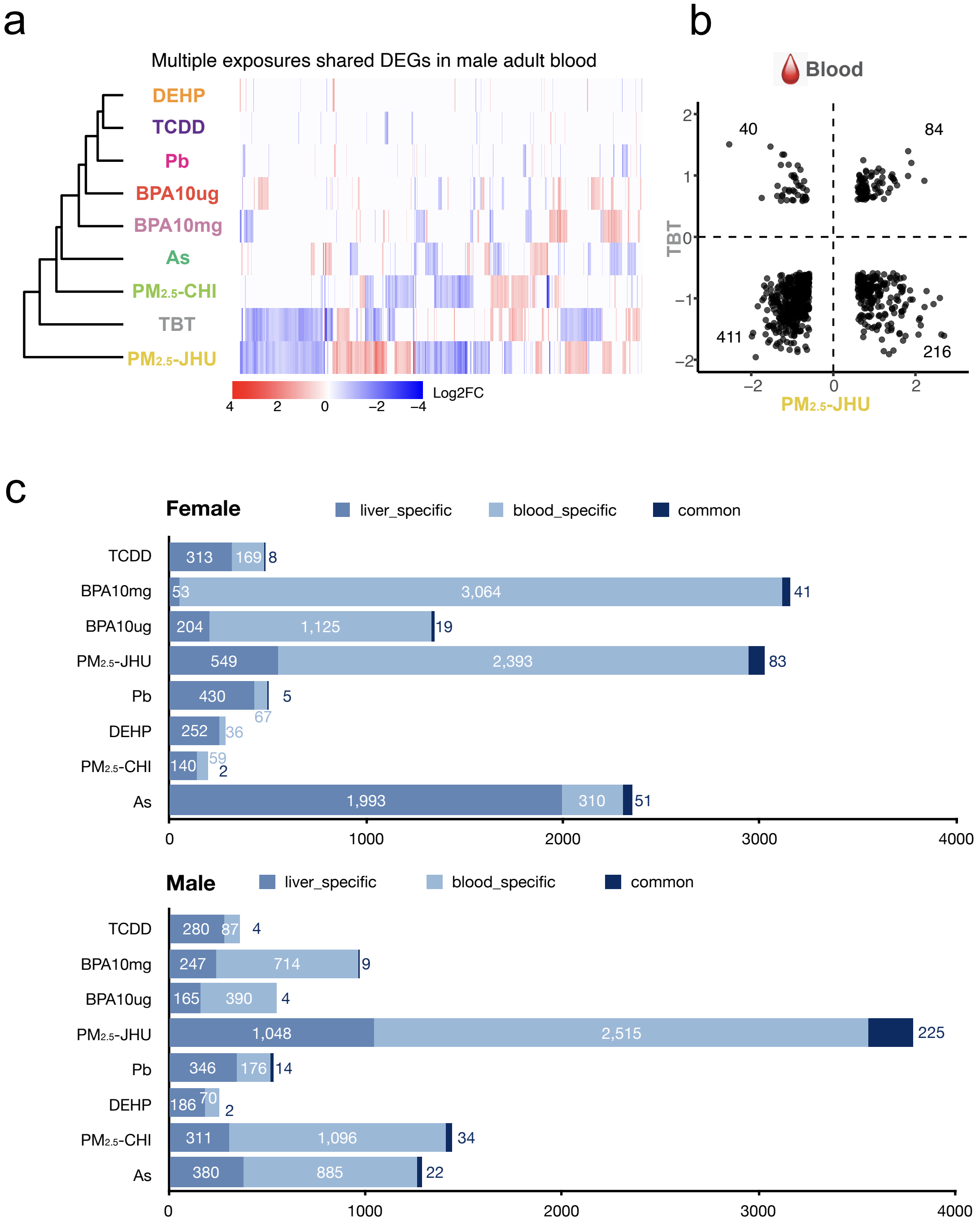
