## Supplementary material for "Comprehensive Transcriptomic and Epigenomic Insights into Environmental Toxicant Exposures: The TaRGET II Resource": S-Figure-3

### Supplementary Figure 3

#### a PCA plot for RNA-seq samples of 5 month blood

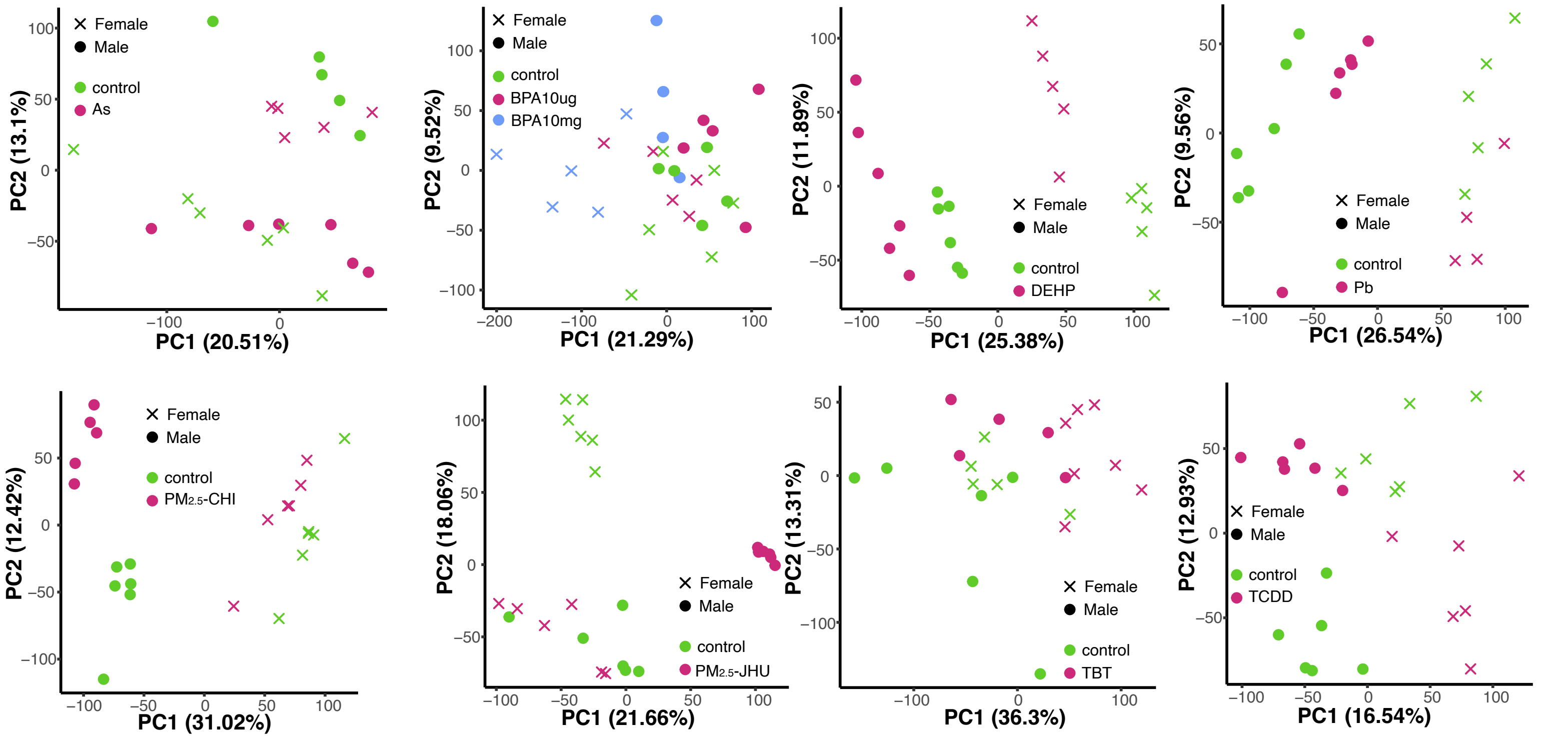


#### b PCA plot for ATAC-seq samples of 5 month blood

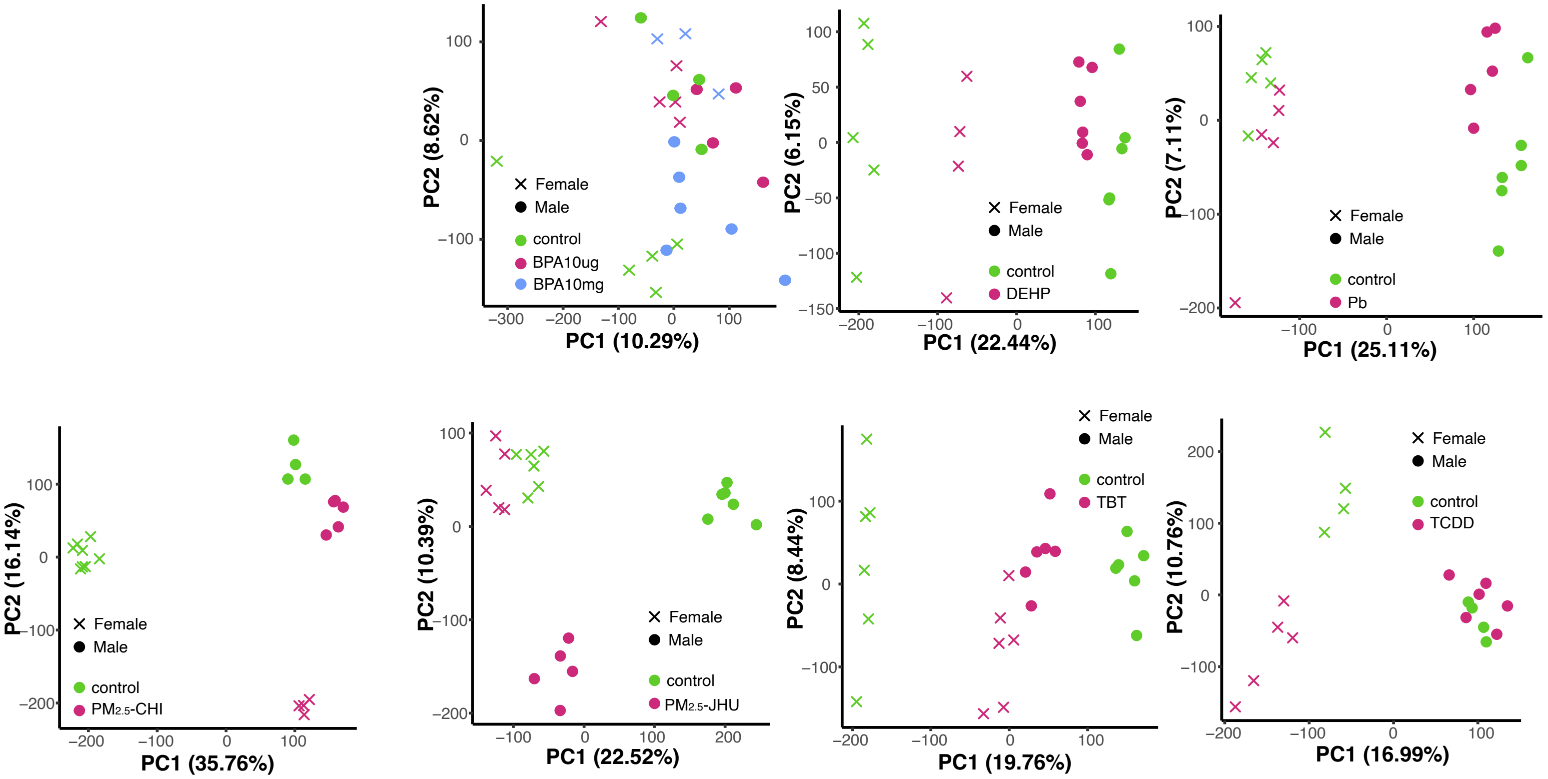
