## Supplementary material for "Comprehensive Transcriptomic and Epigenomic Insights into Environmental Toxicant Exposures: The TaRGET II Resource": Method

**Methods**

**Animal exposure paradigm and tissue collection**

C57BL/6 (B6; Jackson Laboratory, Bar Harbor, ME) mice were used for all experiments, except for the lead (Pb) and phthalate (DEHP) exposure studies, which employed wild-type non-agouti (a/a) mice derived from a >230-generation colony of viable yellow agouti (Avy) mice. These a/a mice are genetically invariant and exhibit 93% genetic identity with the C57BL/6J strain^1^. Mice were exposed to toxicants perinatally via maternal diet, drinking water, or air breathing, with the exposure window spanning from pre-conception through weaning. A schematic overview of the experimental design is provided in **Figure 1**. In brief, two weeks before mating, virgin female dams (6–8 weeks old) were randomly assigned to one of nine exposure groups or a control group:

1) 50 mg/kg diet of BPA (BPA 10mg);

2) 50 µg/kg diet of BPA (BPA 10ug);

3) 25 mg/kg diet of DEHP (DEHP);

4) 32 parts per million (ppm) Pb-acetate drinking water (Pb);

5) PM <2.5 micrometer (PM2.5-JHU);

6) PM <2.5 micrometer (PM2.5-CHI);

7) 0.5 mg/kg BW tributyltin (TBT) drinking water;

8) 1 ug/kg per BW of dioxin (TCDD) via oral gavage;

9) 10 parts per billion (ppb) sodium arsenite drinking water (As).

Female mice (dams) were mated with virgin males (7–9 weeks old) following two weeks of exposure to the assigned experimental diets or water. Exposure continued throughout pregnancy and lactation until offspring were weaned at postnatal day 21 (PND21). Animals were housed in polycarbonate-free cages under a 12-hour light/dark cycle. All experimental procedures were approved by the Institutional Animal Care and Use Committees (IACUC) at each participating production site and were conducted in accordance with the guidelines established by the NIEHS TaRGET II Consortium, adhering to the highest standards of animal welfare^2^.

As, Pb, and TBT exposures were administered via drinking water provided ad libitum. BPA (high and low dose) and DEHP were administered via diet provided ad libitum. TCDD was administered via oral gavage. PM2.5 exposures were administered via a breathing control chamber as scheduled.

**As exposure**

All animal procedures were approved by the University of Pittsburgh Institutional Animal Care and Use Committee. Mice were maintained on an AIN-93G diet (Research Diets, NJ) free of fishmeal and rice products to avoid potential arsenic contamination, and were provided drinking water containing either 0 or 10 parts per billion (ppb) sodium arsenite (≥90% purity; Sigma-Aldrich), refreshed every other day. Both chow and water were batch-tested for arsenic content. Female nulliparous mice were exposed to 0 or 10 ppb inorganic arsenic (iAs) in drinking water beginning two weeks prior to mating and continuing through the end of lactation (eight weeks total). After weaning, F1 offspring were no longer exposed to iAs.

**BPA exposure**

For BPA exposure, a modified AIN-93G diet supplemented with 50 mg/kg BPA (TD.06156; Harlan Teklad) was used for the high-dose group, and a modified AIN-93G diet supplemented with 50 µg/kg BPA (TD.110337; Harlan Teklad) was used for the low-dose group, as previously described^3^. All diet ingredients were sourced from Harlan Laboratories (Madison, WI), with BPA obtained from Sigma-Aldrich (St. Louis, MO). Drinking water was provided in polypropylene bottles. The estimated daily BPA intake was approximately 10 mg/kg body weight (bw) for the high-dose group and 10 µg/kg bw for the low-dose group. Throughout the experiment, animals were maintained on a phytoestrogen-free modified AIN-93G chow (Envigo TD.95092; 7% corn oil diet, Harlan Teklad).

**DEHP exposure**

DEHP was dissolved in corn oil (Envigo) to prepare a stock solution used to generate a customized 7% corn oil chow for experimental exposures. The DEHP concentration was selected to achieve a target maternal dose of 5 mg/kg/day, based on an estimated body weight of 25 g and a daily chow intake of 5 g for pregnant and nursing female mice. This dosing strategy was informed by previous studies demonstrating obesity-related phenotypes following perinatal DEHP exposure at similar levels^4^, and falls within the range of human exposure levels^5^ . Throughout the experiment, animals were maintained on phytoestrogen-free modified AIN-93G chow (Envigo Td.95092, 7% corn oil diet, Harlan Teklad).

**Pb exposure**

Pb-acetate drinking water was prepared by dissolving Pb(II) acetate trihydrate (Sigma-Aldrich) in distilled water to a final concentration of 32 ppm, modeling human-relevant perinatal exposure levels. Previous studies confirmed that this dosing regimen results in maternal blood lead levels (BLLs) ranging from 16 to 60 μg/dL (mean: 32.1 μg/dL)^6^. All exposure water was prepared in a single batch, and Pb concentrations were validated using inductively coupled plasma mass spectrometry (ICP-MS; NSF International; limit of detection = 1.0 µg/L). Animals were maintained on phytoestrogen-free modified AIN-93G chow (Envigo Td.95092, 7% corn oil diet, Harlan Teklad) for the duration of the experiment.

**TBT exposure**

For TBT exposure, animals were administered tributyltin (TBT) chloride (96% purity; Sigma-Aldrich) via drinking water, as previously described^7^. Water consumption was surveyed in non-pregnant females to estimate the administered TBT dose. Consumption was found to increase sharply for approximately three days following parturition, during the onset of lactation. Based on an average intake of 10 mL/day, animals received 200 mL of 3.07 μM TBT solution weekly, corresponding to an estimated dose of 0.5 mg/kg body weight/day. This dose falls below the lowest range of the developmental no observed adverse effect level (NOAEL) (5.8–20 mg/kg body weight) as defined by the Concise International Chemical Assessment Documents (CICADs) for developmental toxicity(Benson, 1999). Animals were maintained on phytoestrogen-free modified AIN-93G chow (Envigo Td.95092, 7% corn oil diet, Harlan Teklad) throughout the study.

**TCDD exposure**

For TCDD exposure, female mice were administered 2,3,7,8-tetrachlorodibenzo-p-dioxin (TCDD; Cambridge Isotopes, cat# ED-901-C) dissolved in corn oil via oral gavage. Dams received three doses of 0.34 µg/kg body weight each, totaling 1 µg/kg body weight across the exposure period from preconception through weaning. Dosing occurred at three time points: two weeks prior to mating, approximately embryonic day 7, and approximately postnatal day 14. Control animals received pure corn oil. Throughout the study, all animals were maintained on phytoestrogen-free modified AIN-93G chow (Envigo Td.95092, 7% corn oil diet, Harlan Teklad).

**PM2.5-JHU exposure**

Animals were exposed to either PM2.5 or filtered air (FA) in dedicated exposure chambers. Exposure occurred 5 days per week (Monday through Friday) from 9:00 AM to 5:00 PM, for a total of 8 hours per day, over a period of 7 weeks. Prior to exposure, all animals were acclimated in the housing facility for one week. Throughout the 7-week exposure period, each group of mice (FA or PM2.5) was rotated between two separate exposure chambers to minimize potential chamber-specific effects. No significant differences in locomotor activity or mobility were observed in any of the mice prior to or during the exposure period, ensuring that baseline physical activity was consistent across groups. Mice were provided with control water and food ad libitum throughout the duration of the PM2.5 exposure.

**PM2.5-CHI exposure**

Dams were placed in either the PM or control chamber for 8 hours daily and mated overnight with male mice until pregnancy was confirmed, as previously described^8^. Upon confirmation of pregnancy, dams continued exposure to either PM2.5 or filtered air for 8 hours per day until the pups were born. Both dams and pups were then exposed to their respective conditions (PM2.5 or filtered air) for 8 hours per day until the pups reached 3 weeks of age.PM2.5 was concentrated from ambient air in Chicago using a chamber connected to the Versatile Aerosol Concentration Enrichment System (VACES), as described previously^9^. The chemical composition of the PM2.5 used in this study has been previously reported^9^. Control mice were exposed to filtered air in an identical chamber connected to the VACES, where a Teflon filter was placed on the inlet valve to remove all particles. Mice were provided with control water and food ad libitum throughout the duration of the PM2.5 exposure.

At postnatal day 21 (PND21), all offspring were weaned and transitioned to control water and control chow. Consortium-wide, one set of male and female mice from each exposure group was euthanized at 3 weeks of age for tissue collection to perform molecular analyses. A second set of males and females were maintained until 5 months of age, and a third set of males and females were maintained until 10 months of age. This study evaluates animals collected at all three time points (3 weeks, 5 months, and 10 months of age).

This study included the following exposure groups: 25 As, 34 BPA high, 33 BPA low, 32 DEHP, 35 Pb, 51 TBT, 24 TCDD, 24 PM2.5-JHU, 46 PM2.5-CHI, and 227 control mice, resulting in a total sample size of 531 animals (Na = 531). Blood, liver, cortex, heart, lung, and other tissues were collected from all animals. All animals and collected tissues were included in subsequent analyses, with no exclusions necessary. For organ and tissue collection, mice were fasted for four hours during the light cycle, beginning in the morning. In the afternoon, animals underwent CO2 euthanasia followed by cardiac puncture, and whole-body perfusion with saline (Sigma). The cortex and liver were then dissected, immediately flash frozen in liquid nitrogen, and stored at -80°C. Tissue collection and processing followed protocols established by the TaRGET II Methods Working Group. Blood was collected in EDTA-coated tubes (Fisher, Cat. #365974), and plasma was separated by centrifugation. Red blood cells were lysed using erythrocyte lysis solution (Qiagen, Cat. #79217), and the remaining white blood cells were washed in PBS and resuspended in Buffer RLT (Qiagen, Cat. #79216) for subsequent DNA extraction. Standardized operating procedures (SOPs) for the consortium’s tissue collection can be accessed on our website at <https://targetepigenomics.org/documents/> , including SOP_Blood_Processing, SOP_Mouse_Liver_Collection, and SOP_Whole_Mouse_Perfusion. For each mouse, one investigator administered the treatment and was therefore aware of the treatment group allocation. All investigators performing subsequent molecular assays were blinded to the treatment group until the treatment groups were revealed during bioinformatics analyses.

**DNA extraction and whole-genome bisulfite sequencing (WGBS)**

DNA extraction was performed using the AllPrep DNA/RNA/miRNA Universal Kit (Qiagen, Cat. #80224). Genomic DNA (gDNA) was used in the preparation of WGBS libraries at the University of Michigan Epigenomics Core. gDNA was quantified using the Qubit BR dsDNA kit (Fisher, Cat. #Q32850), and quality assessed using Agilent’s Genomic DNA Tapestation Kit (Agilent, Cat. #A63880). One WGBS library was prepared for each animal and identified with a unique dual index identifier for deconvolution. For each sample, 200 ng of gDNA was spiked with 0.5% of unmethylated lambda DNA (Cat. no. D1521; Promega) and sheared using a Covaris S220 (10% Duty Factor, 140W Peak Incident Power, 200 Cycle/Burst, 55s). A 2 µl aliquot of processed gDNA was taken to assess shearing using an Agilent High Sensitivity D1000 Kit (Agilent, Cat. #G2991AA). Once shearing was assessed, the remaining gDNA was concentrated using a Qiagen PCR Purification column and processed for end-repair and A-tailing. Ligation of cytosine-methylated adapters was done overnight at 16°C. Following this, ligation products were cleaned using AMPure XP Beads (Fisher, Cat. #NC9933872) before processing for bisulfite conversion using the Zymo EZ DNA Methylation Kit (Zymo, Cat. #D5001), and incubating the DNA according to the following protocol: 55 cycles of 95°C for 30 seconds followed by 55°C for 15 minutes,. After cleanup of the bisulfite converted products, final libraries were amplified over 10 cycles by PCR using KAPA Uracil+ Ready Mix (Fisher, Cat. #501965287) and NEB dual indexing primers. Final libraries were cleaned with AMPure XP beads, concentration assessed using the Qubit BR dsDNA Kit and library size assessed on the Agilent High Sensitivity D1000 Tapestation Kit. Prior to pooling, each library was quantified using KAPA Library Quantification Kit (Fisher, Cat. #501965234). We constructed two pools of 18 libraries, combining a mix of individuals from different litters, sexes, and exposures, to minimize batch effects, and each pool was sequenced on an Illumina NovaSeq6000 S4 200 cycle flow cell (PE-100) at the University of Michigan Advanced Genomics Core. Unless otherwise stated, all enzymes used in library generation were purchased from New England Biolabs. Adapters with universally methylated cytosines were synthesized by Integrated DNA Technologies (IDT).

**RNA extraction and RNA-seq library construction**

Total RNA was isolated via TRIzol Reagent (Thermo Fisher Scientific, 15596026), Phasemaker Tubes (Thermo Fisher Scientific, A33248) and RNA Clean & Concentrator-5 (Zymo Research, R1013). In brief, rat brain tissues were homogenized in 1 ml of TRIzol Reagent, the tissue lysates were transferred to a pre-spined Phasemaker Tube. 0.2 ml of chloroform was added to the tube, the tubes were then shaken for 15 s. The mixture was centrifuged at 16,000 g for 5 min, the top aqueous phase was transferred to a microcentrifuge tube. An equal volume of ethanol was added to the aqueous phase. The RNA purification with DNase treatment was performed following the manual of the RNA Clean & Concentrator kit. Then the Ribosomal RNAs were removed from 500 ng for the total RNA using the NEBNext rRNA Depletion kit (NEB, E6310). Skipping the mRNA isolation part, RNA-seq libraries were then constructed using 10 ng of rRNA-depleted total RNA with Universal Plus mRNA-seq kit (TECAN, 0520-A01) following the kit manual. 2 × 75 bp, 2 × 100 bp, and 2 × 150 bp paired-end sequencing was run for all libraries on the Illumina platforms.

**ATAC-seq library construction**

ATAC-seq was generated using the omni ATAC-seq protocol for frozen tissues^10,11^. In brief, mouse tissues were homogenized in 2 ml of cold 1× homogenization buffer. Nuclei were layered from the tissue lysate with iodixanol solution. 50,000 nuclei were used in the transposition reaction with 100 nM of Transposase (Illumina, 20034197). The ATAC-seq libraries were prepared by amplifying for 9 cycles on a PCR machine with NEBNext High-Fidelity 2× PCR master mix (NEB, M0541). 2 × 75bp paired-end sequencing was run for all libraries on the Illumina platforms.

**ChIP-seq** **library construction**

ChIP-seq was performed as described previously^7^. Briefly, chromatin was cross-linked with formaldehyde, the reaction was stopped with glycine, and the samples were sonicated with a Bioruptor to obtain fragment sizes of 100–300 bp for ChIP-seq. ChIP-validated antibodies used were directed against H3K4me3 (Active Motif #39915), H3K4me1 (Abcam ab8895), H3K27ac (Abcam ab4729), H3K27me3 (Active Motif #61017), and H3K9me3 (Active Motif #61013). Immunoprecipitated DNA was recovered and purified using the QIAquick PCR Purification kit (Qiagen), and DNA concentration was measured using a NanoDrop spectrophotometer. ChIP-sequencing was performed by the University of Houston Core facility. Sequencing libraries were prepared using the QIAseq FX DNA library kit or the QIAseq Ultralow Input Library Kit and the NextSeq 500 platform.

**Infinium Mouse Methylation BeadChip**

1000 ng of genomic DNA isolated from cortex, or 400-1000 ng of genomic DNA isolated from blood, was treated using the EpiTect Bisulfite Kit (Qiagen 59104) according to manufacturer’s instructions with the following modifications: 125 pg of bACE oligo spike-in was included to assess conversion efficiency and DNA was eluted using 20 µl of 0.7 mM Tris-HCl, pH 8.5. Sixty percent of the final elution volume was immediately used for APOBEC3A (A3A) conversion; the remaining 40 percent was stored at -20 ºC. To a 0.2 mL PCR tube, 5 µL of DMSO, 5 µL of reaction buffer (350 mM SPG (molar ratio 2:7:7— succinic acid–sodium dihydrogen phosphate–glycine), pH 5.5 at 25 °C + 1% Tween 20 (v/v)), and bisulfite-treated DNA and H_2_O to 48.5 µL were added. Samples were incubated at 95 ºC for 5 minutes and immediately transferred to a dry ice-cooled PCR-cooler rack (Eppendorf 022510525). While on dry ice, 1.5 µL of 60 mM APOBEC (A3A) enzyme was added. Tubes were quickly spun down and incubated at 37 ºC for 2 hours. Samples were purified using the Oligo Clean and Concentrator Kit (Zymo Research D4061) according to manufacturer’s instructions with the following modification: Samples were eluted using 15 µL of 0.7 mM Tris-HCl, pH 8.5. 1µL was used for conversion efficiency using a custom pyrosequencing assay. The remainder was used for the Infinium Mouse Methylation BeadChip Array (Illumina).

Pyrosequencing: For each sample, 1 µL of bisulfite-converted or A3A-converted DNA was combined with 10 µL Pyromark PCR Mastermix, 2 µL of Coral Load (both components of Qiagen 978705), 1 µL of 10 µM Forward Primer (GCGCGCCAATAAACAACATCAACTCAACAAATA), 1 µL of 10 µM Reverse Primer (5Biosg/AATAAACAACATCAACTCAACAAATA), and 5 µL of H_2_O were combined. PCR conditions were 1 cycle of 95 ºC for 15 minutes and 40 cycles of 95 ºC for 15 seconds, 55 ºC for 30 seconds, 72 ºC for 15 seconds. 10 µL of PCR product was then analyzed on a Qiagen Q96 MD Pyrosequencer according to manufacturer’s instructions using the following sequence to analyze: AGTATCGTATCGATGATCGTGATCGAGTGATCGATGTATCGTAGTCGAT. The oligo spike-in contains artificially generated methylated and hydroxymethylated cytosines. Only samples with levels ≥ 95% or ≤ 5% (depending on the treatment condition) at the 7 analyzed sites were run on the array.

**Raw sequence data and processing**

Total 3,614 genome-wide data sets were gathered from 3-week-old, 20-week-old and 40-week-old mice exposed to pollutants during early development, including arsenic (As), lead (Pb), bisphenol A (BPA), tributyltin (TBT), di-2-ethylhexyl phthalate (DEHP), 2,3,7,8-tetrachlorodibenzo-p-dioxin (TCDD), and particulate matter less than 2.5 µm in diameter (PM2.5). These data sets included 1,043 RNA-seq data, 639 chromatin accessibility data, 659 histone modification data, 522 DNA methylation data (458 WGBS and 71 RRBS), and 744 Infinium Mouse Methylation BeadChip data.

RNA-seq data were processed as in previous studies.  Raw fastaq files of RNA-seq data were processed by Cutadapt^12^ (v1.16; --quality-cutoff=15,10 --minimum-length=36), FastQC (v0.11.7), and STAR^13^ (v2.5.4b; --quantMode TranscriptomeSAM --outWigType bedGraph --outWigNorm RPM) to do the trimming, QC report and mouse genome mapping (mm10) using our own built pipeline, the TaRGET-II-RNA-seq-pipeline (<https://github.com/Zhang-lab/TaRGET-II-RNAseq-pipeline>). Then, gene expressions across normal and exposure samples were calculated by featureCounts^14^ (v1.5.1) within the pipeline based on GENCODE vM20 gene annotation of mouse genome.

Raw fastq of ATAC-seq data were processed by our own built pipeline based on AIAP^15^, the TaRGET-II-ATAC-seq-pipeline (<https://github.com/Zhang-lab/TaRGET-II-ATACseq-pipeline>) that integrated AIAP packages containing optimized QC reports and analysis pipeline with default parameters to generate the open chromatin regions (OCRs). Then, the consensus regions of OCRs across all exposure and control samples were generated with Index (<https://github.com/Altius/Index>) method used for the downstream analysis. The ATAC-seq signals of consensus OCRs were calculated by using the intersectBed method.

The TaRGET-II-WGBS-pipeline (<https://github.com/Zhang-lab/WGBS_analysis>) was built by using the Cutadapt^12^ (v1.16; --quality-cutoff=15,10 --minimum-length=36) and Bismark^16^ (v0.19.0; --bowtie2 -X 1000 --score_min L,0,-0.6 -N 0 --multicore 2 -p 4) to do the trimming and mouse genome mapping. The pipeline also incorporated quality control, generated user-friendly files for computational analysis, and output genome browser tracks for data visualization.

The TaRGET-II-ChIP-pipeline ([https://github.com/Zhang-lab](https://github.com/Zhang-lab/WGBS_analysis)) was used to process ChIP-seq data. Within the pipeline, Cutadapt^12^ (v1.16; --quality-cutoff=15,10 --minimum-length=36), FastQC (v0.11.7), and BWA^17^ (v0.7.12) were used to do the trimming, QC report and mapping. The methylQA^18^ (v0.2.1) was applied to generate read density and mapping statistics, and the macs2^19^ (v2.1.2) was used to call the peaks of ChIP-seq data for downstream analysis.

**Normalization and Batch Effect Correction for RNA-seq and ATAC-seq Data**

RNA-seq and ATAC-seq datasets were normalized to correct for batch effects arising from different data production centers. Samples with the proportion of reads aligned to chrY relative to reads aligned to chrX were used to verify the sex information from metadata. In general, RNA-seq and ATAC-seq data generated within the same production center were analyzed together. Gene counts from RNA-seq were first normalized using the relative log expression (RLE) method implemented in *edgeR*. Subsequently, factor analysis was performed across exposure groups by applying the residuals calculation of the *RUVr* function in *RUVSeq^20^*. To further remove unwanted technical variations (k=3) unrelated to exposures, including biases from tissue dissection and library preparation, a general linear model fitting process was employed as in previous studies^21,22^.

To remove the bias across production centers, the exposed dataset of distinct exposures were first matched to corresponding control samples and performed modified batch-free data correction inspired by ComBat-seq model^23^. In general, a theoretical “batch-free” control data for each sex was defined as pseudo-control Ctrl_P (CP), which represents the estimated empirical distribution of all the control samples with the same variation factors (tissue, age, sex) across distinct production centers. For each sex, the pseudo-control signal $\bar{X_{s}}$ is defined as the mean signal across all control samples of that sex from all centers:

$$\bar{X_{CP}}=\frac{1}{N_{CC}} \sum_{i=1}^{N_{CC}} X_{i,CC}$$

Meanwhile, since the Euclidean distance between the center control (CC) of each production center and CP naturally represents the systematic batch effect of each production center (**Figure: pseudo-control correction**), we transfer the center-specific Euclidean distance as a normalization weight factor (*nw)* to correct control samples and exposed samples based on the calculation of the expression of each gene. The expression of center control (CC) of each production center is defined as

$${Exp}_{{CC}_{i}}={\sum_{m} \left( Exp\_i\_j \right)}/m$$

$Exp\_i\_j$ is the expression of gene *i* in sample *j* out of *m* control samples, and normalization weight factor (*nw*) is defined as the vector from the center control (CC) to pseudo-control Ctrl_P (CP), and the normalized expression of gene *i* (${N\_Exp}_{{ESj}_{i}}$) of exposed sample *j* (*ESj*) are:

$$\vec{nw}= \overset{\to}{CC,CP}=\sqrt[2]{\sum_{n} {({Exp}_{{CC}_{i}}-{Exp}_{{CP}_{i}})}^{2}}$$

$${N\_Exp}_{{ESj}_{i}}=\vec{nw} \cdot{Exp}_{{ESj}_{i}}$$

Pseudo-controls for females and males were generated by calculating the mean signal of all control samples from each sex across production centers. For each center, control samples were then aligned with the respective pseudo-control by scaling their average signal to match the pseudo-control reference, a process conducted separately for each sex. To maintain biological differences between control and exposed samples, all samples from each center were adjusted using the fold changes derived during the alignment process.

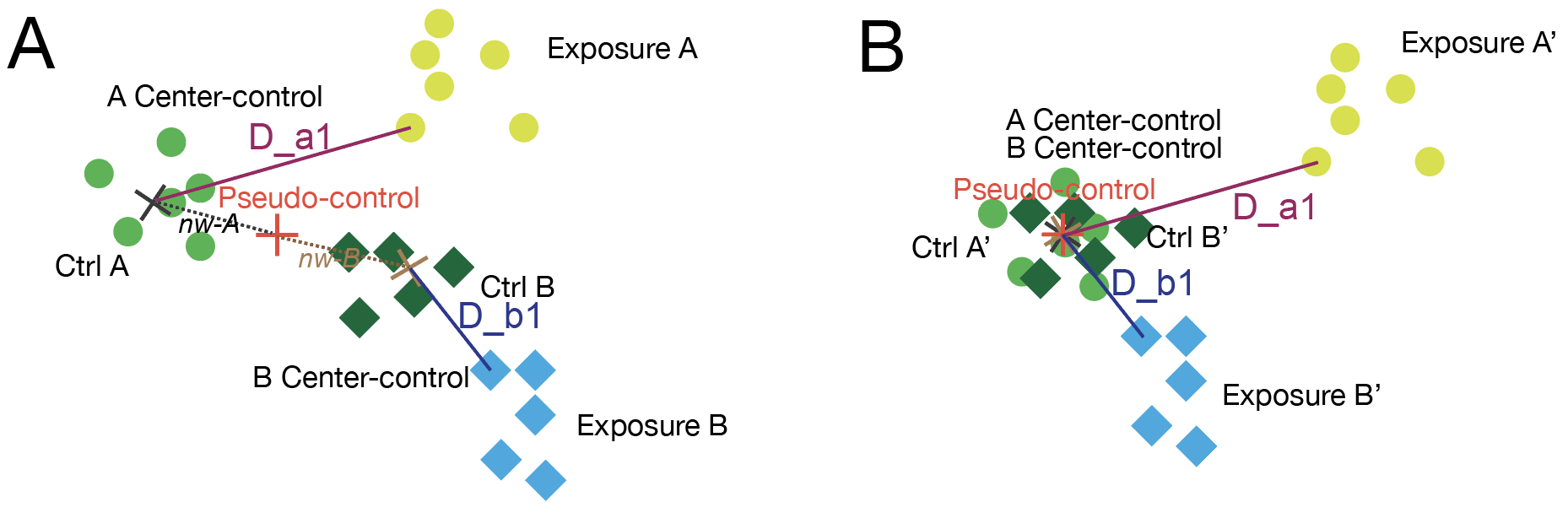

**Figure: pseudo-control correction**. **A).** Data distribution of control samples and exposed samples from two data production cancers in a two-dimensional space. A_Center-control is the centroid of all controls from center A, B_Center-control is the centroid of all controls from center B. D_a1 is the vector from exposed sample a1 and A_Center-control. D_b1 is the vector from the exposed sample b1 and B_Center-control. *nw-A* is the normalization factor for center A samples. *nw-B* is the normalization factor for center B samples. **B).** The samples of center A were normalized by *nw-A,* and the $\vec{D\_a1}$ has remained. Samples of center B were normalized by *nw-B,* and the $\vec{D\_b1}$ has remained.

Read counts of ATAC peaks across all data production centers were normalized in the same way as the RNA-seq data normalization above.

**Differentially analysis**

Differentially expressed genes (DEGs) between control and exposure samples under varying conditions were identified using DESeq2^24^ (v1.34.0) as previous studies^25-27^. Consortium-normalized read count data were used to identify the DEGs between the exposed and control samples. Genes were considered significantly differentially expressed if they met the following stringent criteria: an adjusted p-value (Benjamini-Hochberg correction) less than 0.001 and an absolute log2 fold change greater than log2(1.5), equivalent to a 1.5-fold change in expression.

Differentially accessible regions (DARs) between control and exposure samples were identified using edgeR^28^ (v3.36.0), as previous study^15,29,30^. Consortium-normalized ATAC-seq reads were used to identify DARs, with significance determined by the following cutoffs: an absolute log2 fold change greater than log2(1.5) (a 1.5-fold change in accessibility) and a false discovery rate (FDR) less than 0.01. These criteria balance sensitivity and specificity, ensuring robust detection of chromatin accessibility changes associated with exposure.

Differential methylation regions (DMRs) between control and exposure samples were identified using a method adapted from previous studies^31-34^. Methylation levels were quantified using whole-genome bisulfite sequencing (WGBS) data. Briefly, raw WGBS reads were aligned to the reference genome using Bismark^16^ (v0.23.1) with default parameters, followed by deduplication to remove PCR artifacts. Methylation levels at individual CpG sites were calculated as the ratio of methylated reads to total reads (methylated + unmethylated) at each site, with a minimum coverage of 10 reads per site to ensure reliability. Samples with fewer than 15 million CpG sites or those with less than 60% of CpG sites having a methylation level below 0.7 are excluded from the analyses. Samples with the proportion of reads aligned to chrY relative to reads aligned to chrX greater than 4% were considered male, while those with proportions less than 4% were considered female. If a sample's deduced sex information conflicts with the labeled sex information, it will be excluded from subsequent analyses. For each sample, the mean methylation values for each 1k window across the mm10 genome were calculated and processed using the *normalize.quantiles* function in the preprocessCore R package (v1.60.2). The normalized data were then used in PCA analysis.

DMRs were identified through a genome sliding approach, as previous studies^33-35^: the genome was divided into sliding 200bp windows, the methylated and unmethylated reads counts in each window were calculated in both control samples and exposed samples, and then a Chi-squared test was applied to identify DMRs. Regions were defined as differentially methylated if they exhibited a methylation level difference of at least 0.1 (a 10% absolute difference in methylation between control and exposed condition) and a Q-value (adjusted for multiple testing using the BH method) less than 0.1. Additionally, only regions with at least 2 CpG sites and a minimum average coverage of 20 reads per site were considered to enhance statistical power and reduce noise in the analysis, and the neighboring regions were merged to maximize the width of DMRs.

**Temporal expression patterns across mouse liver development and aging**

The pseudo-control normalized liver transcriptome of control animals and exposed animals was used to explore the developmental expression patterns. The liver transcriptome of the same sex in 3 weeks, 5 months, and 10months were first transferred to normalized z -score to facilitate the downstream calculation:

$$Z_{i,j}=\frac{{(X}_{i,j}-\mu_{j})}{\sigma_{j}}$$

Where $Z_{i,j}$ is the normalized z-score of gene *j* in sample *i*, $X_{i,j}$ is the CPM of gene *j* in sample *i*, $\mu_{j}$ is the mean of expression of gene *j* in all samples, and $\sigma_{j}$ is the standard deviation of the gene *j* in all samples.

The z-scores of all control livers in the same sex were first put into K-means clustering to determine the temporal patterns in three different developmental stages, and the Elbow method of within-cluster sum of squares (WCSS) was used to determine the optimized k, as in previous studies^36-38^.

$$WCSS_{k}= \sum_{j=1}^{k} \sum_{x_{i}\in C_{j}} \left\| x_{i}-\mu_{j} \right\|^{2}$$

Where $C_{j}$ is the *j*-th cluster and $\mu_{j}$ is its centroid. The WCSS of at every *k* was calculated and we noticed that when k=9, the improvement (*I*) of clustering is less than 5%:

$$I_{k}=\frac{(WCSS_{k}-WCSS_{k+1})}{WCSS_{k}}<0.05$$

The calculated $WCSS_{k}$ in both female and male control data were plotted in Supplementary figure 10a, and the optimized k was decided as 9 in both sexes.

To further refine the clustering of temporal gene expression patterns across three life stages (3 weeks, 5 months, and 10 months), an additional Expectation-Maximization (EM) optimization procedure was applied. Initially, temporal expression patterns were defined using K-means clustering. For each cluster, the mean and standard deviation of the z-scores at each time point were used to model the distribution of expression values as a normal distribution. The z-scores of each gene at each stage were then used to calculate the most probable assignment to one of the nine temporal pattern clusters through a PDF function:

$$P_{Gi, Sj}=\frac{1}{\sqrt{{2\pi\sigma}_{h}^{2}}}exp(-\frac{{(z_{Gi}-\mu_{h})}^{2}}{{2\sigma}_{h}^{2}})$$

Where $P_{Gi, Sj}$ is the probability of the gene $i$(expression $z_{Gi}$) in the stage $j$ for the cluster pattern $h\in(1,9)$. Then the probability-likelihood of the gene $i$ belonging to the pattern $h$ is:

$$l_{i, h}=\sum_{i=1}^{n} log\left( \sum_{j=1}^{h} (P_{Gi, Sj}) \right)$$

Gene-to-cluster assignments were updated only if the maximum likelihood of the new assignment improved by more than 5% compared to the previous iteration. Following each reassignment, cluster parameters were recalculated by updating the mean and standard deviation of z-scores based on the newly assigned genes. This iterative EM process was repeated until convergence, defined as no further changes in gene classification across clusters. Gene expression patterns in exposed animals were classified into the best-matched temporal expression clusters using the previously described maximum likelihood-based approach.

**Gene ontology enrichment analysis of DEG**

Biological process enrichment, transcription factor binding, and gene–disease association analyses were performed using the ToppFun^39^ tool from the ToppGene Suite with default parameters, based on gene sets from each temporal expression cluster. Enriched terms were visualized in **Figure 4** using the geom_point function from the ggplot2 R package. The proportions of genes that shifted temporal expression patterns were visualized using the geom_bar function. Alluvial plots depicting changes in temporal expression patterns across time points were generated using the ggalluvial and ggplot2 packages.

**Molecular Responses to Environmental Exposures in Surrogate Blood Cells**

Differentially expressed genes (DEGs), differentially accessible regions (DARs), and differentially methylated regions (DMRs) in female and male blood samples were identified as described above. Visualization of the number of DEGs, DARs, and DMRs across exposures was performed using the geom_bar function from the ggplot2 R package. Shared DEGs and DARs across one or more exposures were identified based on matching gene symbols and genomic coordinates, respectively. Shared DMRs were identified through genomic intersection using the intersectBed function from the bedtools suite. To visualize expression changes of DEGs common to multiple exposures, the pheatmap package was used. DEGs that responded to the same exposure in both blood and liver tissues were identified by matching gene symbols.

**Gene ontology enrichment analysis of DARs**

Enriched biological process terms associated with DARs from different exposure groups were identified using the Genomic Regions Enrichment of Annotations Tool^40^ (GREAT, version 4.0.0) with the following settings: (1) species assembly: Mouse (NCBI Build 38); (2) background regions: whole genome; (3) association rule: Basal plus extension; and (4) significance threshold: Binomial FDR Q-value < 0.05.

**Sex score calculation**

To assess how environmental exposures affect overall gene expression related to sex differences, we defined a 'sex-distance' metric based on the Euclidean distance between pseudo-control female and pseudo-control male samples in three-dimensional principal component (PC) space, a feature-reduced space derived from global gene expression data, as described in previous studies^41,42^. This sex-distance reflects the transcriptomic divergence between sexes and is expected to increase during sex differentiation and morphogenesis, eventually stabilizing after the establishment of sex-specific gene expression patterns post sexual maturation. By comparing exposed samples to this reference distance, we evaluated how different exposures disrupted, attenuated, or exaggerated normal sex-related transcriptional trajectories. The standard sex difference (D) was defined as:

$$D= \sqrt{\sum_{i=1}^{n=3} {(PC.M_{i}-{PC.F}_{i})}^{2}}$$

The sex discrepancy score (S) was defined as the Euclidean distance between a given sample and the pseudo-control female within a stage-matched three-dimensional principal component (PC) space constructed from global gene expression profiles. This metric quantifies the transcriptional divergence of each sample from the female reference, enabling assessment of how exposures influence sex-specific gene expression trajectories. The sex discrepancy score (S) is calculated through the distance $d_{f}$ between the sample $d$ and the pseudo-control female, and the distance $d_{m}$ between the sample $d$ and the pseudo-control male (**Figure sex score**):

$$d_{f}= \sqrt[2]{\sum_{i=1}^{n=3} {(d_{i}-P{C.F}_{i})}^{2}}$$

$$d_{m}= \sqrt[2]{\sum_{i=1}^{n=3} {(d_{i}-P{C.M}_{i})}^{2}}$$

$$S=D\times\cos\angle d\cdot PC.F\cdot PC.M$$

$$\cos\angle d\cdot PC.F\cdot PC.M=\frac{d_{f}^{2}+D^{2}-d_{m}^{2}}{2\times d_{f}\times D}$$

$$S=D\times\frac{d_{f}^{2}+D^{2}-d_{m}^{2}}{2\times d_{f}\times D}$$

In which $d_{i}$ is the value on the principal component $i$ of the given sample $d$, and $P{C.F}_{i}$ is the value on the principal component $i$ of the pseudo-control female, and $P{C.M}_{i}$ is the value on the principal component $i$ of the pseudo-control male.

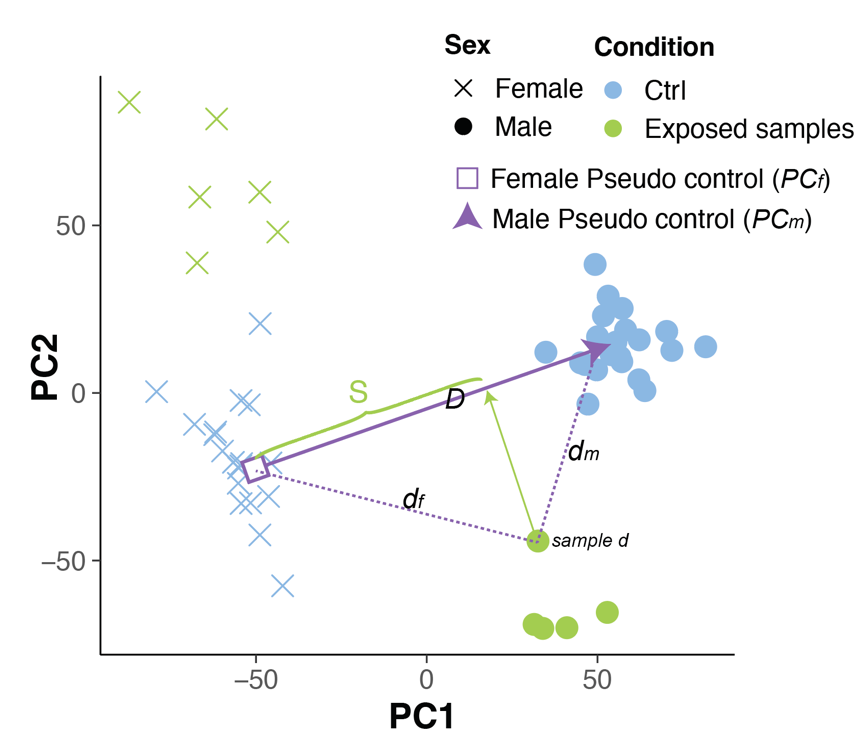

**Figure Sex score**: 2D demonstration of the calculation of sex score (S) for sample *d.* d*_f_* and *d_m_* were the distances from the sample *d* to the pseudo control female and male. *D* is the standard distance between the pseudo control female and male.

The real calculation was made in the 3D PCA space.
